# Transposon accumulation at xenobiotic gene family loci: a comparative genomic analysis in aphids

**DOI:** 10.1101/2023.02.20.529256

**Authors:** Tobias Baril, Adam Pym, Chris Bass, Alex Hayward

## Abstract

The evolution of resistance is a major challenge for the sustainable control of pests and pathogens. Thus, a deeper understanding of the evolutionary and genomic mechanisms underpinning resistance evolution is required to safeguard health and food production. Several studies have implicated transposable elements (TEs) in xenobiotic resistance evolution in insects. However, analyses are generally restricted to one insect species and/or one or a few xenobiotic gene families (XGFs). We examine evidence for TE accumulation at XGFs by performing a comparative genomic analysis across 20 aphid genomes, considering major subsets of XGFs involved in metabolic resistance to insecticides: Cytochrome P450s, glutathione S-transferases, esterases, UDP-glucuronosyltransferases, and ABC transporters. We find that TEs are significantly enriched at XGFs compared to other genes. XGFs show similar levels of TE enrichment to housekeeping genes. But unlike housekeeping genes, XGFs are not constitutively expressed in germline cells, supporting the selective enrichment of TEs at XGFs, rather than enrichment due to chromatin availability. Hotspots of extreme TE enrichment occur around certain XGFs. In aphids of agricultural importance, we find particular enrichment of TEs around cytochrome P450 genes with known functions in the detoxification of synthetic insecticides. Our results provide evidence supporting a general role for TEs as a source of genomic variation at host XGFs, and highlight the existence of considerable variability in TE content across XGFs and host species. These findings demonstrate the need for detailed functional verification analyses to clarify the significance of individual TE insertions and elucidate underlying mechanisms at TE-XGF hotspots.

## Introduction

Xenobiotics are substances that are foreign to an organism, including naturally occurring plant allelochemicals and man-made insecticides (Li et al. 2007). Xenobiotic toxicity can present a strong selective pressure, leading to the rapid evolution of resistance via efficient xenobiotic removal, metabolism, and tolerance (Chung et al. 2007; Li et al. 2007). Consequently, numerous insects have evolved resistance to both plant allelochemicals, produced for plant defence, and synthetic insecticides, developed to protect economically important crops from insect pests, or humans and livestock from insect vectored diseases (Sparks et al. 2021; Sternberg and Thomas 2018). The intensive use of numerous synthetic insecticides has led many insects to evolve resistance to multiple insecticide classes (Li et al. 2007; Bradshaw et al. 2016), severely affecting our ability to control target insects (Singh et al. 2021). The emergence of multiple insecticide resistance is a major global challenge, threatening global food security in the case of agricultural pests, and human and animal health in the case of disease vectors (Worner and Gevrey 2006; Deutsch et al. 2018).

Two major mechanisms are frequently implicated in conferring xenobiotic resistance across a wide range of arthropods (Li et al. 2007; Kliot and Ghanim 2012): (i) ‘target-site resistance’, involving structural changes (mutations) in the gene encoding the insecticide target protein, that make it less sensitive to the toxic effect of the insecticide; (ii) ‘metabolic resistance’, involving the increased production or activity of metabolic enzymes that break down or sequester the insecticide. Regarding the latter, several major gene families are associated with metabolic resistance, which act during three distinct phases of xenobiotic metabolism (Kennedy and Tierney 2013): (i) Cytochrome P450 monooxygenases (CYP genes); (ii) glutathione S-transferases (GSTs), esterases, and UDP-glucuronosyltransferases (UGTs); (iii) ATP-binding cassette transporters (ABC transporters). We briefly summarise these three phases below. During phase I, CYP genes catalyse functionalisation reactions, where hydrophobic xenobiotics are typically converted to more water-soluble metabolites through the addition of functional groups (Kennedy and Tierney 2013). CYP genes are diverse enzymes involved in several purposes from biosynthesis to metabolism, and are sometimes referred to as ‘nature’s blowtorch’, due to their high-valence iron chemistry oxidation mechanism (Guengerich 2009). CYP genes can mediate resistance to all classes of insecticides due to their broad substrate specificity and versatility, whilst they are also involved in several other processes, such as juvenile hormone and ecdysteroid metabolism, and fatty acid biosynthesis (Feyereisen 2005; Kennedy and Tierney 2013; Feyereisen 1999). During phase II, GSTs, UGTs, and esterases catalyse conjugation reactions between phase I substrates and endogenous molecules to form water-soluble metabolites (Kennedy and Tierney 2013). During phase III, ABC transporters, which are cell membrane transport proteins that efflux toxins and modified toxins from the cell, export xenobiotics and products of phase I and II (Gott et al. 2017). In addition, ABC transporters can also block the cellular import of xenobiotics to protect the host, which is termed phase 0. (Gott et al. 2017). Insecticide resistance can arise via mutations acting on the genes involved in the processes described above through various genetic mechanisms, including up-regulation, changes to coding sequence, and gene amplification (Li et al. 2007). It is also important to note that many xenobiotic gene family (XGF) members are involved in other metabolic processes beyond xenobiotic resistance. For example, some GSTs have roles in processes including eye pigmentation, intracellular transport, and cell signalling pathways (Ketterman et al. 2011; Ranson and Hemingway 2005), whilst ABC transporters have numerous roles in processes including heme biosynthesis, iron homeostasis, and protection against oxidative stress (Dermauw and Van Leeuwen 2014). UGTs also have roles in the modulation of endobiotics and olfactory processes (Ahn et al. 2012), and esterases have roles in neurodevelopment and pheromone signalling (Gilbert and Gill 2014).

There is growing evidence that transposable elements (TEs) can play an important role in the evolution of xenobiotic resistance in insects (Gilbert et al. 2021; Rostant et al. 2012), with examples from several major lineages: Diptera: *Drosophila* (Feyereisen 2005; Bogwitz et al. 2005; Schmidt et al. 2010; Aminetzach et al. 2005; Remnant et al. 2013; Mateo et al. 2014; Salces-Ortiz et al. 2020), *Culex* (Itokawa et al. 2011; Darboux et al. 2007)*, Musca domestica* (Li et al. 2007; Kasai and Scott 2001), *Anopheles funestus* (Weedall et al. 2020) Lepidoptera: *Heliothis* (Gahan et al. 2001; Yang et al. 2007; Zhao et al. 2010; Chen and Li 2007), *Pectinophora gossypiella* (Fabrick et al. 2011; Wang et al. 2019) Hemiptera: *Myzus persicae* (Singh et al. 2020; Panini et al. 2021). TEs are DNA sequences capable of moving from one genomic location to another within the genome. TEs occur in nearly all eukaryotic genomes, and are implicated in the evolution of host genomic novelty through diverse processes including the modification of regulatory networks, chromosomal rearrangements, exon shuffling, and donation of coding sequence (Bourque et al. 2018; Wells and Feschotte 2020; Cosby et al. 2019; Sundaram et al. 2014). In the case of xenobiotic resistance, TEs are reported to have contributed to evolvability via myriad mechanisms, including: gene amplification and duplication (Singh et al. 2020; Remnant et al. 2013), knockout of a susceptible allele in heterozygous individuals (Panini et al. 2021), increases in detoxification gene expression (Bogwitz et al. 2005; Itokawa et al. 2011), allelic succession leading to increases in resistance gene copy number (Schmidt et al. 2010), and alternative splicing and production of truncated proteins that prevent interactions with xenobiotics (Darboux et al. 2007; Gahan et al. 2001; Yang et al. 2007; Zhao et al. 2010).

Studies examining the contribution of TEs to resistance evolution suggest that there can be an additive effect of successive TE insertions at focal host loci over time. For example, in the well-known case of the cytochrome P450 (CYP) gene *CYP6G1* and DDT resistance in *Drosophila*, successive TE insertions are linked with an increasing ability to detoxify the insecticide (Catania et al. 2004; Daborn et al. 2002; Chung et al. 2007; Schmidt et al. 2010). A similar process involving the accumulation of TEs, in this case at *CYP6CY3,* is implicated in the evolution of resistance to nicotine, following a host plant shift to tobacco in the aphid *Myzus persicae nicotianae* (Puinean et al. 2010; de Little et al. 2017). More widely, there is evidence that the selective accumulation of TEs at rapidly evolving host loci under strong selection may represent a general evolutionary process. For example, studies have reported the enrichment of TEs at other gene classes, such as immune genes and those involved in responses to external stimuli (van de Lagemaat et al. 2003; Song et al. 1998). Existing studies strongly suggest that TEs are involved in individual cases of resistance to certain insecticides, but there is currently very limited understanding of how associations with TEs vary among different classes of XGFs, or among whole clades of insects. However, patterns in TE accumulation remain poorly described, and it is unclear to what extent TEs may be enriched at XGFs compared to other host genes, or how much variability exists across XGFs.

Here we consider patterns of TE accumulation at XGFs in the genomes of 20 species sampled from across aphid phylogeny, for which high-quality genome assemblies are available. The aphid family is a relevant clade within which to explore the genomic bases of insecticide resistance, as it includes numerous globally important crop pests responsible for causing billions of US dollars of crop losses annually (Van Emden and Harrington 2017), and members that have been intensively targeted with insecticides and evolved resistance. For example, the severely damaging global crop pest *Myzus persicae* (the green peach aphid) is resistant to at least 82 insecticides, via at least eight different resistance mechanisms (Bass et al. 2014; Mota-Sanchez and Wise 2023; Singh et al. 2021). We hypothesise similar broadscale patterns for TE accumulation at XGFs among aphid genomes, given similar challenges from xenobiotics that target conserved biological pathways. Meanwhile, given the strong selection pressure that can arise from xenobiotic exposure, and repeated xenobiotic challenges over time, we anticipate an ability to detect patterns at the species-level.

Since the TE landscapes of aphids are poorly described, we begin by characterising patterns of TE content across our focal genomes. After this, we examine key fundamental questions relating to the pattern of associations between TEs and XGFs in aphid genomes. (i) Firstly, we test whether XGFs are enriched for TEs compared to other host genes, to evaluate the overall signature of TE accumulation at XGFs. (ii) Secondly, we examine the extent to which TE content varies among individual XGFs and among major XGF classifications, to determine if contributions from TEs are more pronounced at certain types of XGF. (iii) Thirdly, we consider if specific TE classifications are enriched at XGFs (e.g DNA TEs, rolling circles, *Penelope*-like elements, LINEs, SINEs, and LTRs), to explore if particular TEs are predisposed to contribute to the evolution of host resistance. (iv) Fourthly, we assess whether patterns of TE accumulation at XGFs vary among aphid species to examine if patterns vary across aphid phylogeny. (v) Fifthly, we investigate TE enrichment at XGFs that have demonstrated associations with the detoxification of synthetic insecticides, to test if these show particularly strong signatures of TE recruitment. (vi) Lastly, we explore whether our results suggest a selective role for patterns in TE accumulation at XGFs versus the alternative explanation of increased availability for TE insertion due to chromatin availability.

## Results

### Transposable Element Landscapes in Aphids

TEs make considerable contributions to genome size in eukaryotes (Kidwell 2002; Gregory 2005). In aphids, we find that TE content varies from 8.21% (*Melanaphis sacchari*) to 35.52% (*Metopolophium dirhodum*) of total genome size (Figure 1B, Supplemental Table S1). This is in accordance with levels of TE content identified among a large-scale survey of 196 insect species, where TE content varied from 0.08% to 53.93% of total genome size (Peccoud et al. 2017). Similarly, consistent with findings in other eukaryotic lineages, we report a significant strong positive association between genome size and TE content in aphids (Linear regression, F_1,19_=46.35, p<0.01, R^2^=0.69, Supplemental Figure S1). Aphids show variation in sexual and asexual lifecycles (Van Emden and Harrington 2017). It is hypothesised that asexual lineages would suffer from Muller’s ratchet as deleterious mutations accumulate (Muller 1964). Despite this, we find no significant difference in TE content between holocyclic and anholocyclic lineages (Wilcoxon Rank Sum, W_14,6_ = 35, p>0.05) (Figure 1B).

**Figure 1.**
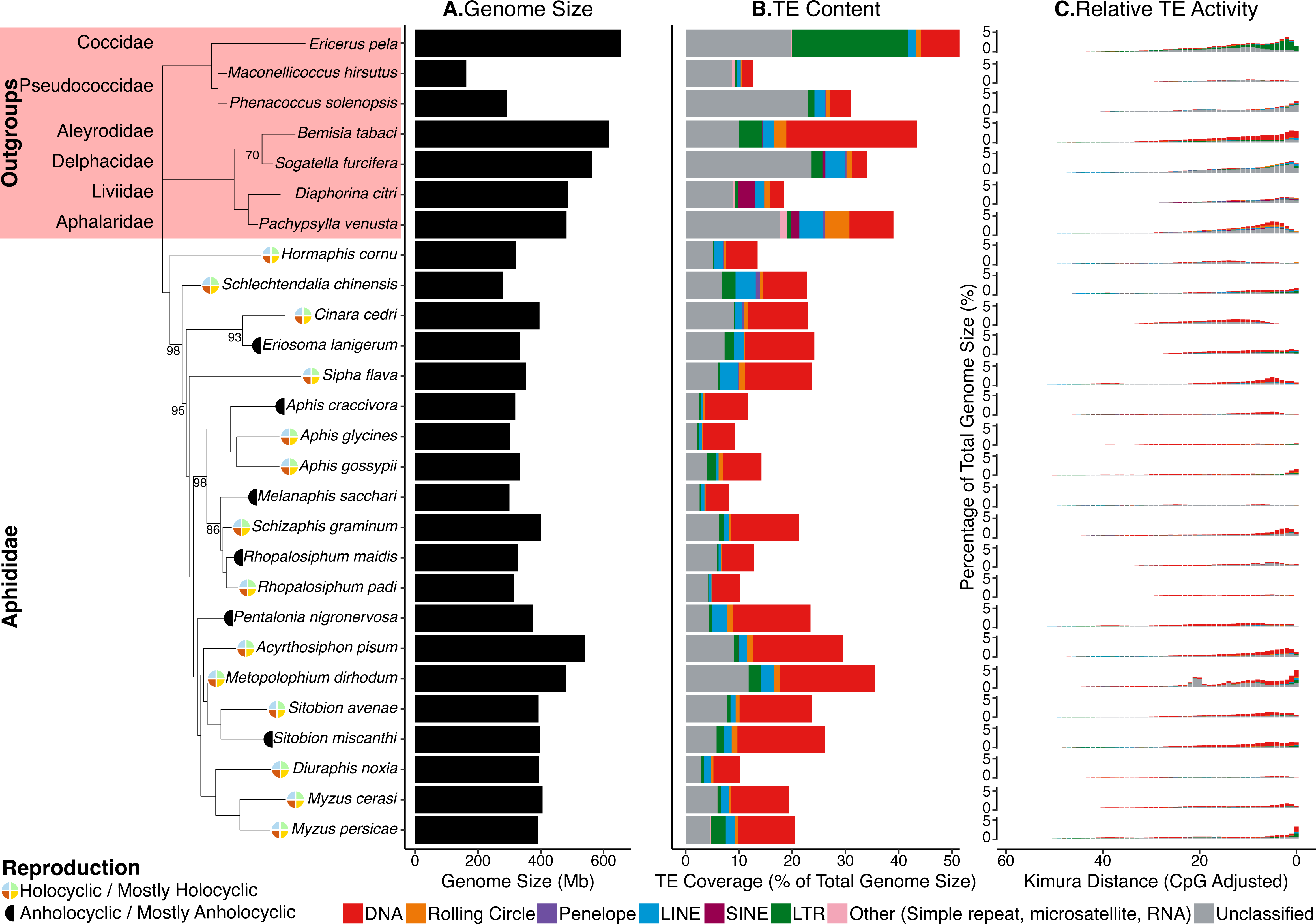
Summary of TE content and activity in aphid genomes, plotted in phylogenetic order. Nodes in the aphid phylogeny with bootstrap support <100 are labelled. Reproductive mode (Van Emden and Harrington 2017) is labelled next to species names. Genome size is represented by the black bars. TE content is expressed as percentage of total genome size for each species, with major TE classifications represented by the colours indicated in the key. Kimura 2-parameter distance (CpG adjusted) from each TE family consensus is used as a proxy for relative TE activity, where a lower Kimura distance indicates more recent TE activity. Due to challenges with accurately estimating TE age, divergence from consensus better reflects relative TE activity, with lower Kimura distances signalling more recent TE activity. Activity is organised such that recent TE activity is towards the RHS of the X axis.

TEs have a non-random distribution within host genomes (Bourque et al. 2018). As expected, given the predominantly deleterious effects of TE insertion into host genes (Bourque et al. 2018; Sultana et al. 2017), exonic regions exhibit the lowest levels of TE sequence content in aphid genomes (mean TE content in exons = 1.90% of total genome size). Across 19 of the 20 aphid genomes considered here, the greatest TE coverage (expressed as proportion of total genome size) is found in 5’ and 3’ gene flanking regions, where it varies between 3.49% (*M. sacchari*) and 17.94% (*M. dirhodum*) (Figure 2B). However, this pattern is not universal across all aphid species. In *Sipha flava,* the majority of TE coverage is present in intronic regions (10.60%), while in *M. cerasi*, the majority occurs in intergenic space (11.28%) (Figure 2B).

**Figure 2.**
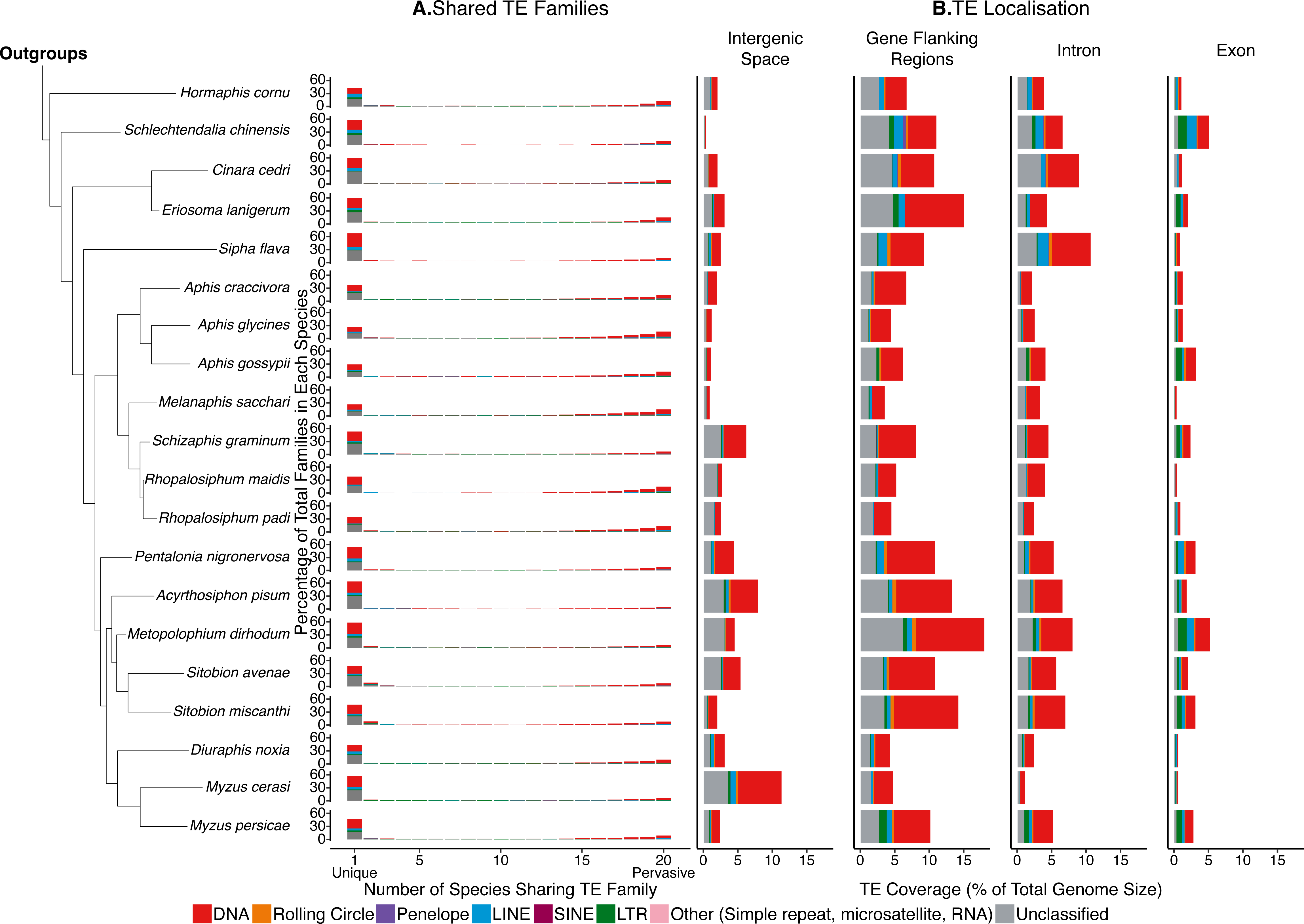
Overview of the extent to which TE families are shared among aphid genomes, and their insertion locations. A. Quantification of shared and unique TE families in aphids. Main TE classifications are represented by the colours indicated in the key. B. TE genome compartment occupancy, expressed by TEs as a percentage of total genome size. Gene flanking regions are defined as 20kb directly upstream or downstream of the gene body.

The amount of TE sequence and the frequency of TE insertions in genic versus intergenic regions in aphid genomes is not significantly correlated with genome compactness, described as the ratio of genic (intron and exon) to intergenic (gene flanks and intergenic space) base pairs (Spearman’s Rank, S=1680, p>0.05, rho=-0.09, Supplemental Figure S2, Supplemental Table S2). *S. flava* has the most compact genome, with a genic:intergenic base pair ratio of 1.50:1 (250.0 Mb:166.8 Mb), and so presumably a higher likelihood of TE insertion into genic regions compared to intergenic space, leading to the observation that the majority of TE insertions are found in intronic regions. In contrast, *M. cerasi* has the least compact genome, with a much lower genic:intergenic base pair ratio of 0.13:1 (76.1 Mb:559.2 Mb). In less compact genomes such as *M. cerasi*, TEs can accumulate in expanded intergenic regions that act as ‘genomic safe havens’, where insertion is less likely to exert deleterious effects (Arkhipova 2018). For context, the human genome (GCF_009914755.1), has a TE content of 44%-69% (Nurk et al. 2022; de Koning et al. 2011) and a genic:intergenic base pair ratio of 0.55:1 (1717.7 Mb:3117.3 Mb), making it several times more compact than *M. cerasi*, but much less compact than *S. flava*.

Across all aphid species and in most individual genomes, the dominant TE classification is DNA TEs, which comprise between 36.6% (*Schlechtendalia chinensis*) and 68.9% (*Aphis craccivora*) of total TE content (μ = 45.27%, SD = 18.00%, Figure 1B, Table 1, Supplemental Table S1). SINEs are present only in the genomes of 7 of the 20 species considered (Supplemental Table S1) and, where present, they comprise just 0.009% (*A. pisum*) to 0.1% (*A. craccivora*) of total TE content.

**Table 1.**
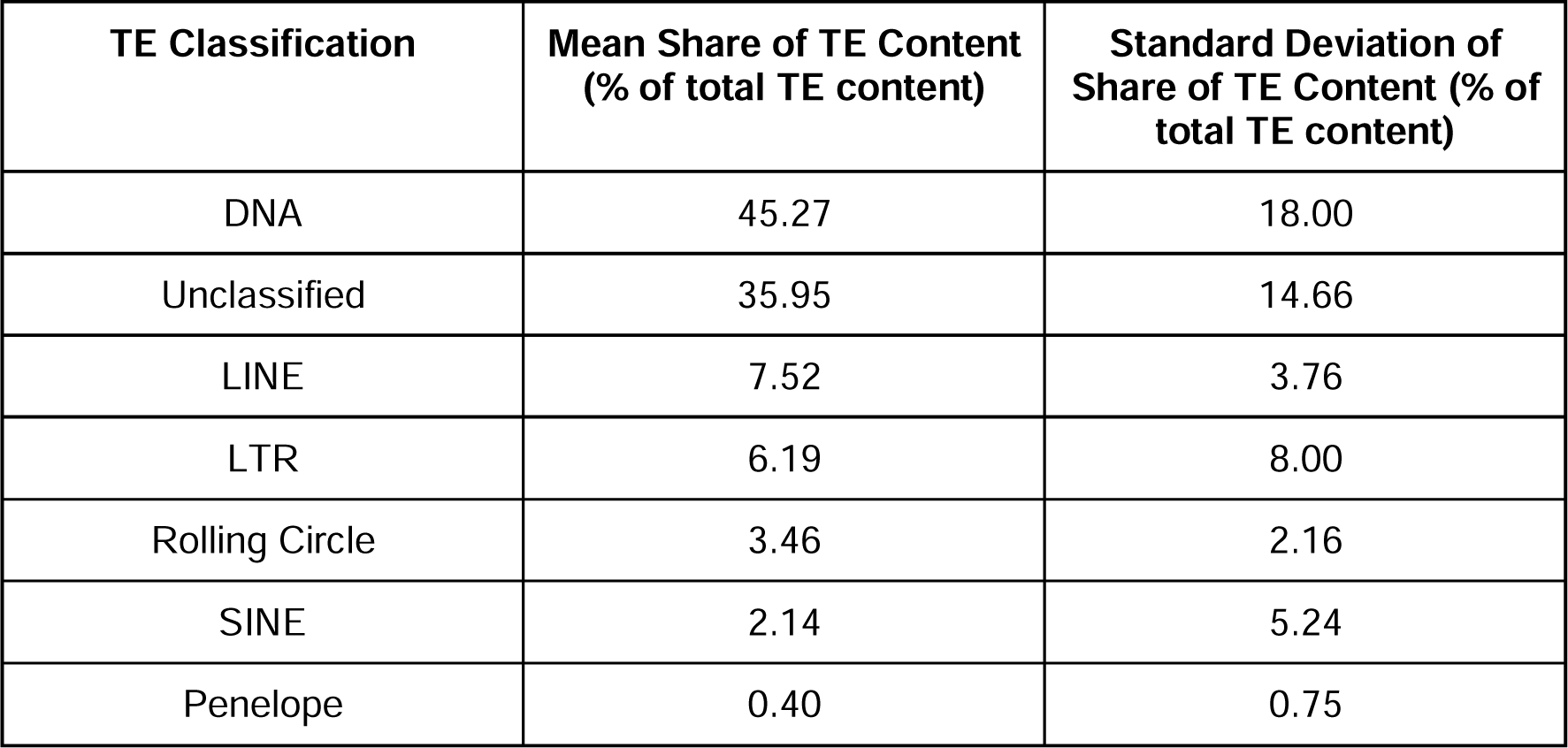
Mean and standard deviation of the percentage of total TE content attributed to each of the main TE classifications for the 20 aphid genomes included in this study.

There is little evidence of ancient TE activity in aphid genomes, as indicated by the relatively low number of TEs showing high divergence from their respective TE consensus models in repeat landscape plots (Figure 1C). On the other hand, there is considerable evidence of relatively recent TE activity in most aphid species, with particular evidence of very recent TE activity in *Myzus persicae* and *M. dirhodum*, indicated by large numbers of TEs with low genetic distance from their respective consensus sequences (Figure 1C). The relative absence of ancient TE activity is presumably a consequence of a relatively rapid genomic turnover rate in aphids, and contrasts strongly with TE landscapes in mammals, where a low rate of genomic turnover leads to considerable accumulations of ageing TEs (Blass et al. 2012). Instead, the dynamics we observe for aphids are consistent with patterns reported for other insects, such as lepidopterans (Lavoie et al. 2013; Baril and Hayward 2022).

The extent to which TE content differs among the genomes of closely related species can vary greatly. We find no significant effect of phylogeny on TE abundance (No. of TEs present in a genome) or TE diversity (No. distinct TE families present in a genome (Wicker et al. 2007)) in aphids (Phylogenetic GLMM, TE abundance: posterior mean = 0.592, 95% HPD = 0.005,0.999; TE diversity: posterior mean = 0.022, 95% HPD = 0.000,0.111). Thus, more closely related aphid species do not possess similar numbers of TEs or a similar level of TE diversity. We also find that most TE families present in aphid genomes are species-specific, with almost half of the TE families present in a particular aphid genome being unique to that species (μ = 46.97%) (Figure 2A). On average, 10.02% of the TE families identified in an aphid genome are shared among all 20 species (Figure 2A). The remaining 43.01% of TE families are shared in similar proportions among 2 to 19 aphid species (i.e. μ = 2.53% per category, Figure 2A). The strong signature of species-specificity observed for aphid TE families suggests that the independent gain of new TE families is a major factor driving aphid TE landscapes. Collectively, these findings suggest a dynamic repeat landscape in aphid genomes, characterised by relatively frequent gain and turnover of TEs.

### TEs are enriched at XGFs compared to other host genes

Previous studies have implicated TEs in the evolution of xenobiotic resistance across a range of insects and resistance loci (Li et al. 2007). However, the extent to which TE association at XGFs represents a general mechanism for the evolution of new resistance phenotypes remains unclear.

Across all aphid genomes considered here, we find significant enrichment of TEs, both in TE coverage and TE count, at XGFs in comparison to all other genes excluding XGFs (TE coverage: Wilcoxon Rank Sum, W_6488,401787_ = 732,129,770, p<0.01; TE count: Wilcoxon Rank Sum, W_6488,401787_ = 770,552,556, p<0.01) (Figure 3, Supplemental Figure S3). Specifically, XGFs in aphids have a mean of 2.4x TE coverage and 2.1x TE count compared to all other genes.

**Figure 3.**
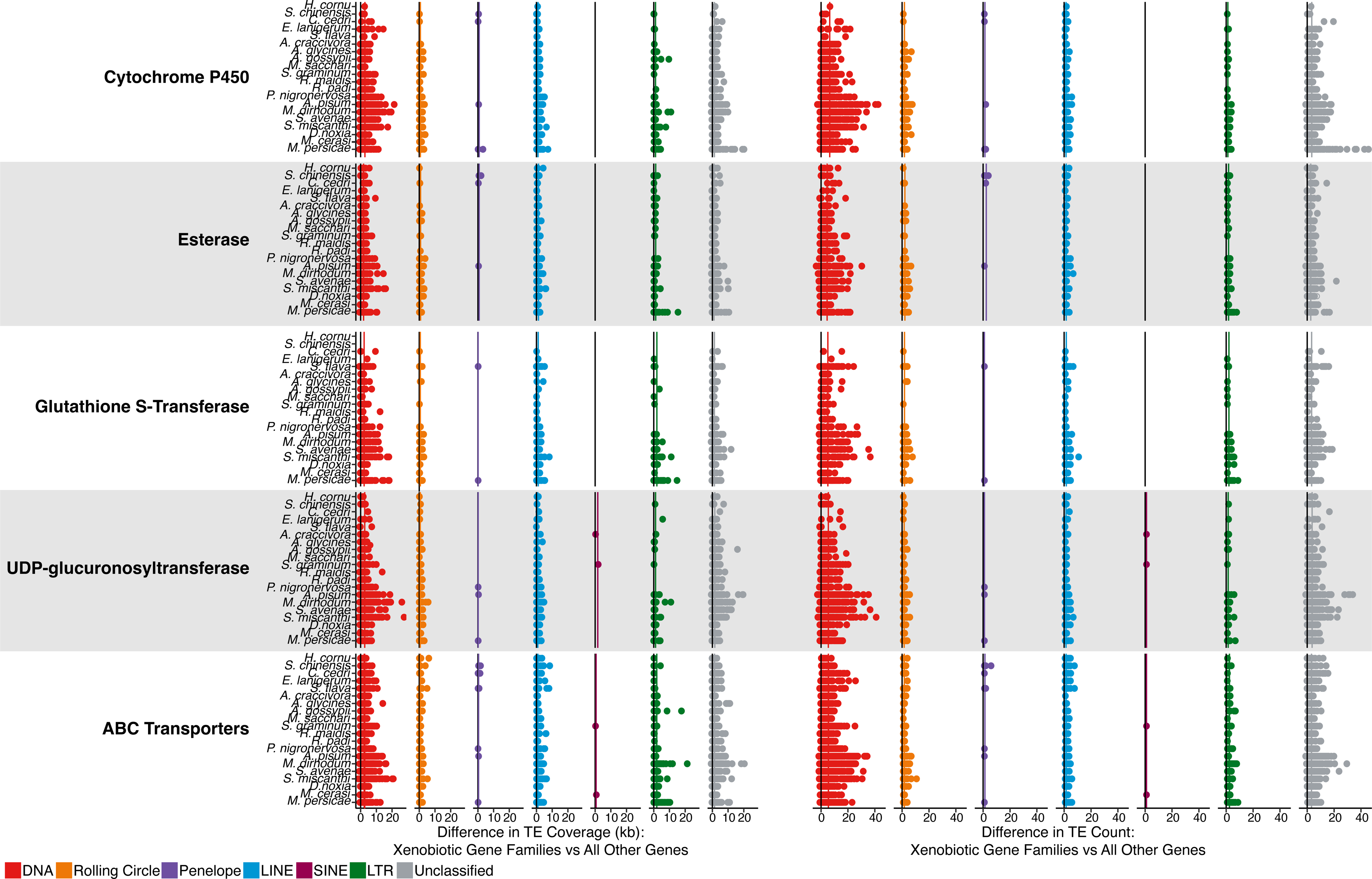
Differences in TE coverage and TE count at XGFs compared to all other host genes. Coverage and count of genic (exon and intron) XGF sequence plus flanking regions (20kb directly upstream or downstream of the gene body) is shown. Each point represents a single XGF locus. Black lines indicate the expected TE coverage difference whereby XGFs and other genes are equally enriched for TEs (i.e. 0 enrichment). Coloured lines indicate species mean coverage and count for each XGF locus and TE type. Major TE classifications are indicated by the colours in the key.

### TE content varies among individual XGF loci

We observe an enrichment of TEs around all major XGF categories considered compared to other host genes: Cytochrome P450s (CYPs), 2.4x TE coverage, 2.1x TE count; glutathione S-transferases (GSTs), 2.7x TE coverage, 2.3x TE count; esterases, 2.0x TE coverage, 1.8x TE count; UDP-glucuronosyltransferases (UGTs), 2.0x TE coverage, 1.7x TE count; ABC transporters (ABCs), 2.8x TE coverage, 2.4x TE count (Figure 3, Supplemental Figure S3, Supplemental Table S3). Whilst all major XGF categories are enriched for TEs, there is no significant difference in relative enrichment among XGF types, suggesting relatively equal levels of enrichment across XGF types (Kruskal-Wallis, TE sequence: ^2^ = 7.72, p = 0.10; TE count: χ^2^_4_ = 4.25, p = 0.37). However, considerable variability in TE content is present within XGF categories, attributable to extremely large accumulations of TEs at certain individual XGFs (Table 2, Figure 4). Some of these XGFs are unusually large in terms of sequence length compared to the mean size for the XGF type in question, as indicated by gene size *Z*-scores > 3 (Table 2), which is potentially a consequence of the increased presence of TEs in their intronic regions (Figure 4). To investigate this, we consider gene sizes with genic TEs removed. We find that 5/18 of the most TE-enriched XGFs have significantly inflated gene sizes when TE insertions are removed, indicated by a *Z*-score of > 3. This suggests that these loci were significantly larger than average before accounting for TE contributions. However, the remaining 13 highly TE-enriched XGFs have gene sizes within the expected range in the absence of TEs, suggesting that TEs are responsible for the significant increases in gene size at the majority of TE-enriched XGF loci. Individual XGFs with the greatest enrichment of TE coverage and TE count are listed in Table 2 (i.e. >35kb TE sequence and >80 TE insertions, compared to a non-XGF mean of 1.5kb TE sequence and 1.48 TE insertions). Whilst all XGF types are represented in the TE hotspots listed in Table 2, there are markedly more hotspots at UGTs and CYP genes, and fewer at ABCs, GSTs, and Esterases.

**Figure 4.**
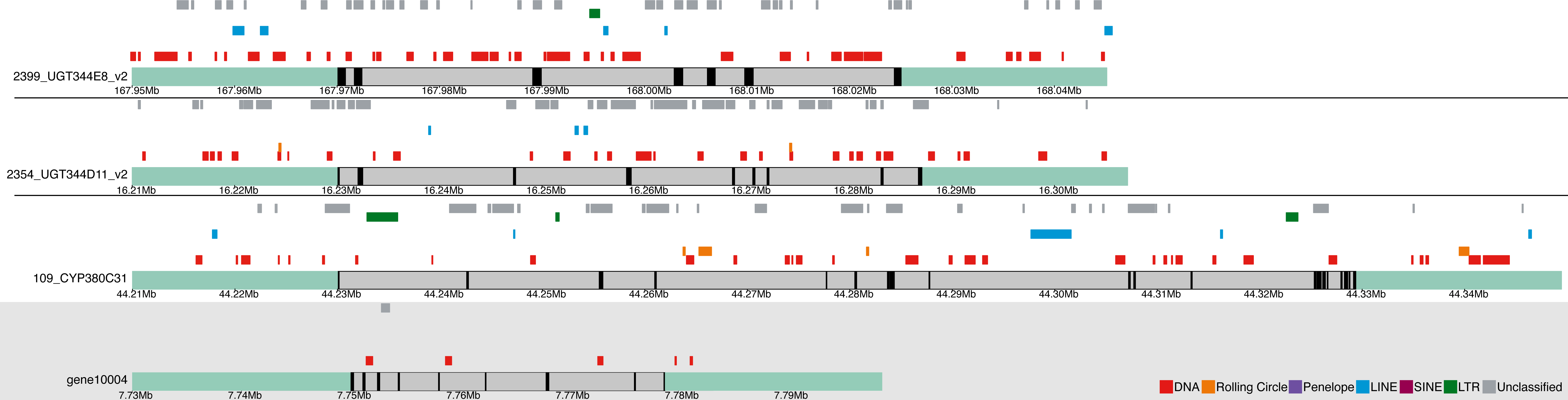
Karyoplots illustrating the most TE-dense XGF loci, corresponding to the XGFs highlighted in bold in Table 2. For comparison, the gene region represented in the shaded box (gene10004) shows a non-XGF from M. persicae with representative TE coverage close to the mean for non-XGFs (1,519bp). XGFs 125_CYP380C52 and 105_CYP380C19 are not shown, as these are nested at the CYP cluster containing 109_CYP380C31. Sea green shaded regions indicate gene flanking regions. Grey bars represent gene bodies, and black bars show exonic regions. TEs are annotated above their respective gene tracks, with colours indicating the main TE classifications, as depicted in the key at the bottom of the figure.

**Table 2.** XGF loci with the greatest enrichment for TE coverage and TE count in aphids. *Z*-scores indicate the number of standard deviations away from mean gene size for individual genes. *Z*-scores <-3 or >3 indicate statistically unusually small or large genes compared to mean XGF size, respectively. Entries in bold are XGFs that are highly enriched for both TE coverage and TE count. ABC transporter subfamilies have been labelled where appropriate (Dermauw and Van Leeuwen 2014).

| Aphid Species | Gene ID | XGF Type | Mean Gene Size (by gene type) | Gene Size in bp (Z-score) | XGF associated TE Coverage (bp) | Fold-increase above aphid-wide enrichment (XGF-type specific) | Fold-increase above aphid-wide enrichment (non-XGF genes) | XGF associated TE Count | Fold-increase above aphid-wide enrichment (XGF-type specific) | Fold-increase above aphid-wide enrichment (non-XGF genes) | Gene Size after removing genic TEs (Z-score) |
| --- | --- | --- | --- | --- | --- | --- | --- | --- | --- | --- | --- |
| Greatest Enrichment of TE Coverage |  |  |  |  |  |  |  |  |  |  |  |
| *Acyrtosiphon pisum* | 2399_UGT344E8_v2 | UDP-glucuronosyltransferase | 8,302 | 55,119 (4.63) | 55,257 | 9.05 | 36.38 | 118 | 9.08 | 79.73 | 24,991 (2.45) |
| *Acyrtosiphon pisum* | 2354_UGT344D11_v2 | UDP-glucuronosyltransferase | 8,302 | 57,143 (4.83) | 45,208 | 7.40 | 29.76 | 96 | 7.38 | 64.86 | 26,620 (2.68) |
| *Myzus persicae* | 125_CYP380C52 | Cytochrome P450 | 7,221 | 100,885 (9.70) | 44,205 | 6.28 | 29.10 | 102 | 6.00 | 68.92 | 56,825 (5.97) |
| *Myzus persicae* | 109_CYP380C31 | Cytochrome P450 | 7,221 | 99,455 (9.55) | 42,790 | 6.08 | 28.17 | 94 | 5.53 | 63.51 | 57,937 (6.10) |
| *Myzus persicae* | 352_gst_g525.t1 | Glutathione S-Transferase | 3,450 | 16,231 (2.65) | 40,940 | 5.02 | 26.95 | 45 | 2.50 | 30.41 | 10,854 (1.54) |
| *Myzus persicae* | 105_CYP380C19 | Cytochrome P450 | 7,221 | 61,287 (5.60) | 39,133 | 5.56 | 25.76 | 100 | 5.88 | 67.57 | 34,033 (3.17) |
| *Metopolophium dirhodum* | 2360_UGT344A13 | UDP-glucuronosyltransferase | 8,302 | 18,807 (1.04) | 38,003 | 6.22 | 25.02 | 45 | 3.46 | 30.41 | 8,327 (0.11) |
| *Metopolophium dirhodum* | 1793_UGT345A3 | UDP-glucuronosyltransferase | 8,302 | 6,504 (-0.18) | 37,746 | 6.18 | 24.85 | 29 | 2.23 | 19.59 | 4,684 (-0.40) |
| *Sitobion miscanthi* | 690_UGT341A5_v2 | UDP-glucuronosyltransferase | 8,302 | 4,437 (-0.38) | 36,666 | 6.00 | 24.14 | 42 | 3.23 | 28.38 | 3,950 (-0.50) |
| *Sitobion miscanthi* | 687_UGT341A5 | UDP-glucuronosyltransferase | 8,302 | 4,456 (-0.38) | 36,179 | 5.92 | 23.82 | 39 | 3.00 | 26.35 | 3,969 (-0.50) |
| *Metopolophium dirhodum* | 13_abc_g2222.t1 | ABC Transporter (ABC Subfamily G) | 11,211 | 14,244 (0.32) | 35,841 | 4.30 | 23.60 | 35 | 1.84 | 23.65 | 12,513 (0.06) |
| *Sitobion miscanthi* | 1630_UGT344E8_v2 | UDP-glucuronosyltransferase | 8,302 | 3,249 (-0.50) | 35,637 | 5.84 | 23.46 | 57 | 4.38 | 38.51 | 2,150 (-0.75) |
| *Sitobion miscanthi* | 72_est_g6279.t1 | Esterase | 10,888 | 35,579 (1.82) | 35,569 | 5.90 | 23.42 | 63 | 4.50 | 42.57 | 23,910 (1.03) |
| **Greatest Enrichment of TE Count** |  |  |  |  |  |  |  |  |  |  |  |
| *Acyrtosiphon pisum* | 2399_UGT344E8_v2 | UDP-glucuronosyltransferase | 8,302 | 55,119 (4.63) | 55,257 | 9.05 | 36.38 | 118 | 9.08 | 79.73 | 24,991 (2.45) |
| *Myzus persicae* | 125_CYP380C52 | Cytochrome P450 | 7,221 | 100,885 (9.70) | 44,205 | 6.28 | 29.10 | 102 | 6.00 | 68.92 | 56,825 (5.97) |
| *Myzus persicae* | 105_CYP380C19 | Cytochrome P450 | 7,221 | 61,287 (5.60) | 39,133 | 5.56 | 25.76 | 100 | 5.88 | 67.57 | 34,033 (3.17) |
| *Acyrtosiphon pisum* | 2354_UGT344D11_v2 | UDP-glucuronosyltransferase | 8,302 | 57,143 (4.83) | 45,208 | 7.40 | 29.76 | 96 | 7.38 | 64.86 | 26,620 (2.68) |
| *Myzus persicae* | 109_CYP380C31 | Cytochrome P450 | 7,221 | 99,455 (9.55) | 42,790 | 6.08 | 28.17 | 94 | 5.53 | 63.51 | 57,937 (6.10) |
| *Acyrtosiphon pisum* | 2423_abc_g11808.t1 | ABC Transporter (ABC Subfamily A) | 11,211 | 25,566 (1.53) | 25,867 | 3.11 | 17.03 | 84 | 4.42 | 56.76 | 19,631 (0.83) |
| *Sitobion avenae* | 1033_UGT344A27 | UDP-glucuronosyltransferase | 8,302 | 62,310 (5.34) | 33,066 | 5.41 | 21.77 | 82 | 6.31 | 55.41 | 45,645 (5.35) |
| *Acyrtosiphon pisum* | 1088_CYP380C8 | Cytochrome P450 | 7,221 | 10,900 (0.38) | 28,620 | 4.07 | 18.84 | 82 | 4.82 | 55.41 | 5,975 (-0.27) |
| *Sitobion miscanthi* | 911_UGT344A27 | UDP-glucuronosyltransferase | 8,302 | 64,596 (5.57) | 27,402 | 4.49 | 18.04 | 81 | 6.23 | 54.73 | 38,974 (4.41) |
| *Acyrtosiphon pisum* | 1_abc_g11808.t1 | ABC Transporter (ABC Subfamily A) | 11,211 | 39,350 (3.00) | 31,743 | 3.81 | 20.90 | 81 | 4.26 | 54.73 | 25,849 (1.50) |

### Specific TE types are enriched at XGFs

While most TE types contribute to enrichment at XGFs, overall enrichment is primarily driven by increases in DNA TE content, due to the high frequency of DNA TEs in aphid genomes, where they comprise almost half of total TE content (Table 3).

**Table 3.** Mean base pair coverage of each TE classification at XGFs among aphids. Numbers in brackets indicate mean TE copy number at each XGF for each TE classification.

| TE Type | Non-XGF | All XGFs | CYPs | GSTs | Esterases | UGTs | ABCs |
| --- | --- | --- | --- | --- | --- | --- | --- |
| **DNA** | 1,974 (2) | 4,026(10) | 4,445 (11) | 4,800 (11) | 3,513 (9) | 3,743 (8) | 4,609 (11) |
| **Rolling Circle** | 728 (1) | 794 (2) | 836 (2) | 849 (2) | 1,126 (3) | 706 (2) | 756 (2) |
| **Penelope** | 608 (1) | 456 (1) | 840 (2) | 175 (1) | 848 (2) | 185 (1) | 477 (1) |
| **LINE** | 959 (1) | 992 (2) | 965 (2) | 1,225 (2) | 1,492 (2) | 859 (2) | 1,126 (2) |
| **SINE** | 358 (1) | 1,144 (1) | - | - | - | 1,686 (1) | 602 (1) |
| **LTR** | 1,238 (2) | 1,817 (3) | 1,115 (1) | 2,528 (3) | 1,397 (2) | 1,124(2) | 3,826 (5) |
| **Unclassified** | 1,307 (1) | 2,255 (5) | 2,114 (5) | 2,290 (6) | 1,754 (4) | 2,508 (6) | 2,384 (6) |

The TE type with the greatest degree of enrichment in coverage around XGFs in comparison to other genes is SINEs, although there is a slight non-significant reduction in SINE count at XGFs (**TE coverage**, 3.2x enrichment: XGFs μ = 1,144bp, all other genes μ = 358bp; Wilcoxon Rank Sum, W_9,482_ = 439, p < 0.01; **TE count,** 0.82x enrichment: XGFs μ = 1.00, all other genes μ = 1.22; Wilcoxon Rank Sum, W_9,482_ = 2592, p = 0.14). This is notable, since SINEs are generally very scarce in aphid genomes, and suggests a particular retention of SINE sequence at XGFs (Figure 1B, Figure 3, Table 1, Supplemental Table S1). However, given their scarcity, the relative contribution that SINEs make to TE enrichment at XGFs compared to other TE types is extremely low. Conversely, DNA TEs are the most abundant TE type in aphid genomes, and they are significantly enriched at XGFs both in TE coverage and TE count (**TE coverage**, 2.0x enrichment: XGFs μ = 4,026bp, all other genes μ = 1,974bp; Wilcoxon Rank Sum, W_6156,336473_ = 616,739,338, p < 0.01; **TE count**, 6.4x enrichment: XGFs μ = 10.05, all other genes μ = 1.57; Wilcoxon Rank Sum, W_6156,336473_ = 1,267,581,920, p < 0.01) (Figure 3). Rolling circle TEs show significant, but lower levels of enrichment (**TE coverage**, 1.1x enrichment: XGFs μ = 794bp, all other genes μ = 728bp; Wilcoxon Rank Sum, W_1801,55720_ = 46,195,509, p<0.01; **TE count**, 1.8x enrichment: XGFs μ = 2.14, all other genes μ = 1.22; Wilcoxon Rank Sum, W_1801,55720_ = 56,336,944, p < 0.01). In contrast, LINEs do not show significant levels of enrichment at XGFs in terms of TE coverage, but there is a significant enrichment of LINE TE count (**TE coverage**, 1.03x enrichment: XGFs μ = 992bp, all other genes μ = 959bp; Wilcoxon Rank Sum, W_2575,101686_ = 128,128,748, p = 0.06; **TE count**, 1.3x enrichment: XGFs μ = 1.79, all other genes μ = 1.41; Wilcoxon Rank Sum, W_2575,101686_ = 167,286,786, p < 0.01). Similarly, we find only a slight, non-significant, enrichment of LTR TE coverage at XGFs, but we do identify a significant enrichment of LTR TE count (**TE coverage**, 1.4x enrichment: XGFs μ = 1,817bp, all other genes μ = 1,238bp; Wilcoxon Rank Sum, W_1004,51486_ = 25,656,118, p = 0.69; **TE count**, 1.7x enrichment: XGFs μ = 2.54, all other genes μ = 1.51; Wilcoxon Rank Sum, W_1004,51486_ = 32,328,754, p < 0.01). Only PLEs show a depletion, albeit insignificant, in TE coverage around XGFs, but they also show significant enrichment in count around XGFs (**TE coverage**, 0.7x enrichment: XGF μ = 456bp, all other genes μ = 608bp; W_47,6823_ = 185,654, p = 0.06; **TE count**, 1.3x enrichment: XGF μ = 1.43, all other genes μ = 1.11; Wilcoxon Rank Sum, W_47,6823_ = 174,958, p < 0.05). However, like SINEs, PLEs account for a very small proportion of total TE content (Supplemental Table S1).

### XGF enrichment varies among aphid genomes

We find no significant difference in TE enrichment among different XGF types (i.e CYP genes, esterases, UGTs, GSTs, ABCs) (Kruskal-Wallis, TE coverage: χ^2^_4_ = 7.72, p = 0.10; TE count: χ^2^_4_ = 4.25, p = 0.37, Figure 3). However, we do find significant variation in the magnitude of TE enrichment at XGFs among aphid species (linear regression, TE coverage: F_19,78_ = 37.49, p < 0.01, adjusted R^2^ = 0.88, TE count: F_19,78_ = 28.28, p < 0.01, adjusted R^2^ = 0.84). Specifically, in terms of TE coverage, *C. cedri* shows the highest levels of enrichment across all XGFs, with a mean enrichment of 15.9x compared to non-XGFs (Figure 5, Supplemental Table S3). This is considerably higher than the lowest level of mean TE coverage enrichment, which was found in *M. sacchari* with 1.9x TE coverage enrichment at XGFs compared to non-XGFs (Figure 5, Supplemental Table S3). In terms of TE count, the highest level of enrichment at XGFs compared to non-XGFs was also found in *C. cedri*, with a mean TE count enrichment of 28.5x the number of TE insertions, whilst the lowest level was found in *A. craccivora*, with a mean enrichment of 4.6x the number of TE insertions at XGFs compared to non-XGFs (Figure 5, Supplemental Table S3)

**Figure 5.**
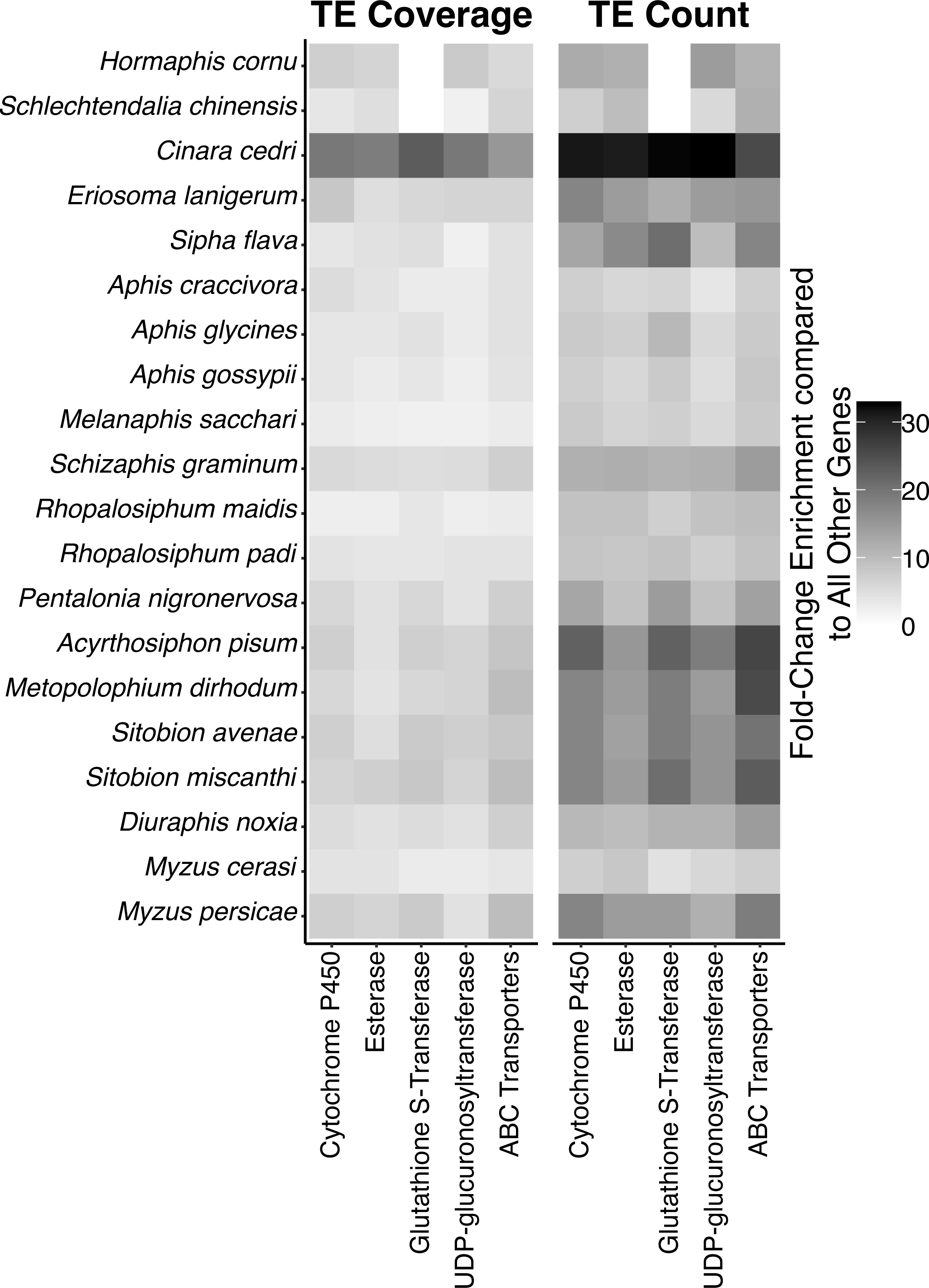
TE enrichment at XGFs compared to genome-wide levels. Fold-change enrichment for TE coverage and TE count at XGFs compared to within-genome levels at non-XGFs across 20 aphid species.

Overall, whilst we observe enrichment in both TE coverage and TE count at XGFs compared to non-XGFs in aphid genomes, fold-change increases are larger for TE count. High enrichment for TE count combined with lower increases in TE coverage suggest that enrichment is driven by the presence of large numbers of TE fragments at XGFs, as opposed to full-length TEs (Figure 6). Indeed, mean TE length at XGFs across aphids is just 536bp, which is much shorter than most full length TEs, excluding SINEs (Wells and Feschotte 2020; Wicker et al. 2007). Further, 90.40% of TEs across XGFs are below 1,000bp in length (Figure 6). Therefore, it is likely that a general pattern in TE accumulation at XGFs is TE truncation with retention of specific TE domains, as previously observed for certain individual cases of TE co-option during the evolution of host insecticide resistance (Singh et al. 2020; Remnant et al. 2013; Panini et al. 2021; Bogwitz et al. 2005; Itokawa et al. 2011; Schmidt et al. 2010; Darboux et al. 2007; Gahan et al. 2001; Yang et al. 2007; Zhao et al. 2010).

**Figure 6.**
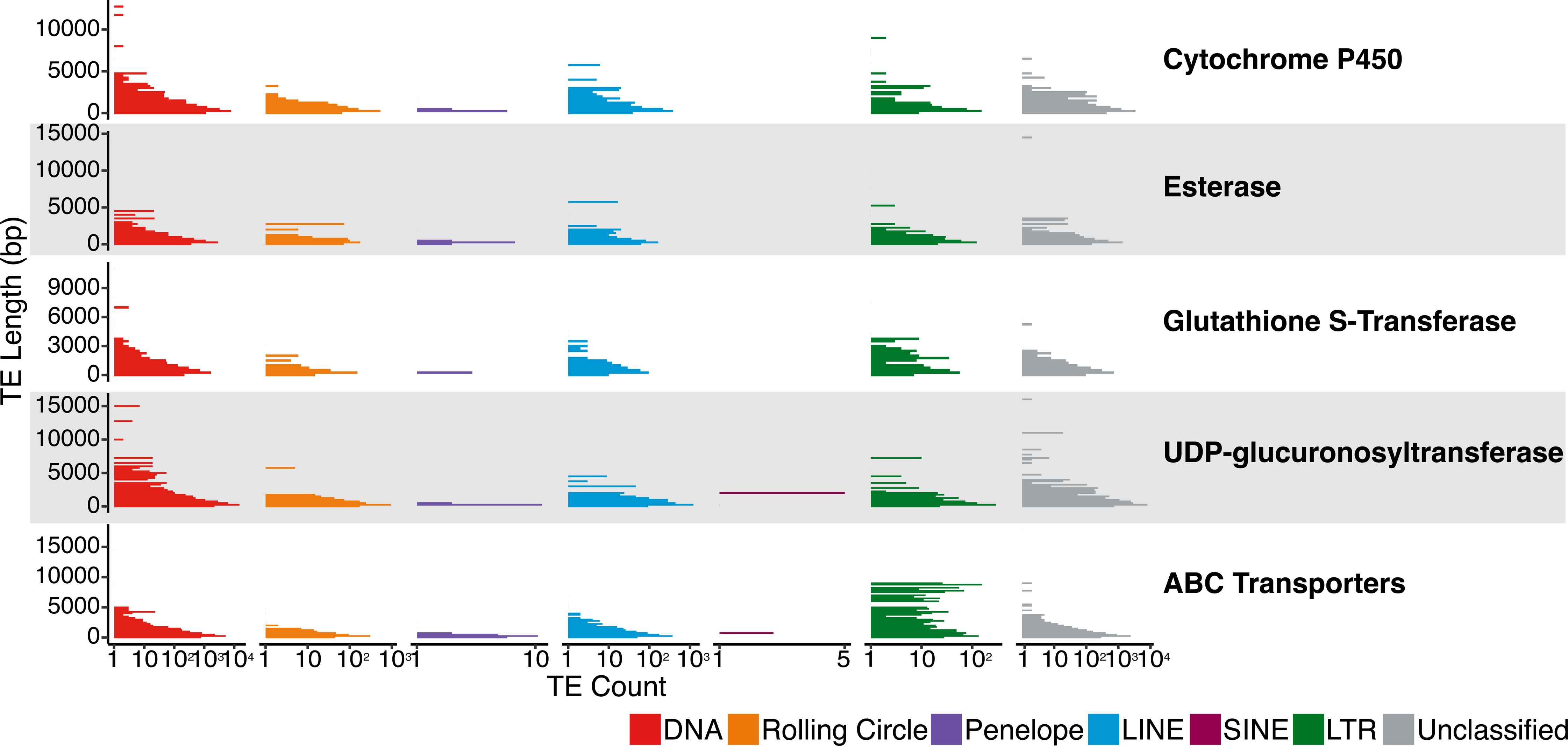
Fragmented TEs at XGFs across all aphid species, presented as TE count by TE length with a bin width of 250bp. TE count is presented on a log10 scale. The main TE classifications are indicated by the colours indicated in the key. Mean TE length across all TEs at XGFs is 536bp.

### TEs are enriched at XGFs associated with the detoxification of synthetic insecticides in certain aphid species

Cytochrome P450 genes (CYP genes) are a highly important gene family for the detoxification of xenobiotics, including insecticides (Katsavou et al. 2022; Feyereisen 2005). Indeed, increases in the expression of certain CYP genes can provide increases in insecticide resistance (Nauen et al. 2022). Additionally, several cases of TE interactions with CYP genes have been characterised (Chung et al. 2007; Chen and Li 2007; Daborn et al. 2002; Itokawa et al. 2011; Schmidt et al. 2010; Karunker et al. 2008; Nauen et al. 2022). Therefore, we undertook a targeted analysis to examine evidence for the accumulation of TEs at CYP genes with putative functions in xenobiotic resistance. For this, aphid CYP genes were labelled based on the availability of evidence to support their xenobiotic resistance functions from (Katsavou et al. 2022), as some CYP gene family members play other roles, such as hormone breakdown. This resulted in the assignment of 2-29 CYP genes per aphid genome to the xenobiotic detoxification category (Supplemental Table S5). In *S. flava*, *C. cedri*, and *M. cerasi*, we find significant enrichment of TEs surrounding CYP genes in CYP gene (sub)families frequently implicated in the breakdown of xenobiotics compared to those with other functions (Figure 7A). However, we find the opposite pattern in *S. graminum*, *M. dirhodum*, and *M. persicae*, where overall there is a significant depletion of TEs surrounding CYP genes associated with xenobiotic resistance (Figure 7A).

**Figure 7.**
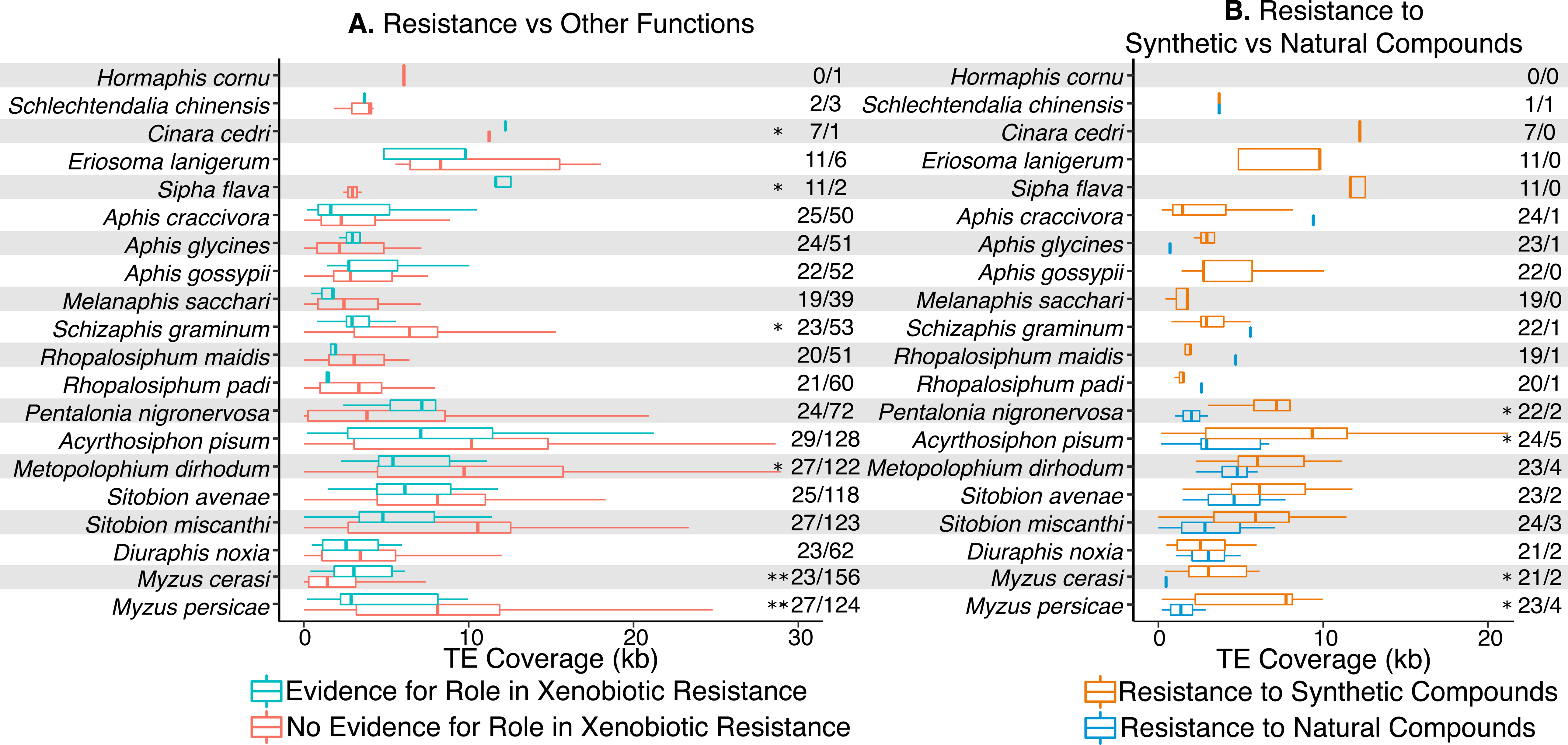
TE association with cytochrome P450 genes in aphids. A. TE coverage around CYP genes linked to xenobiotic resistance versus those with other functions. Numbers on the right indicate the number of genes in each group being compared. B.TE coverage around CYP genes linked to resistance to synthetic insecticides versus those conferring resistance to natural xenobiotics. Numbers on the right indicate the number of genes in each group under comparison.

Next, we considered patterns of TE accumulation at CYP genes specifically involved in resistance to synthetic insecticides, rather than resistance to xenobiotics more generally. For this, CYP genes associated with xenobiotic resistance functions were labelled to reflect the compounds to which they confer resistance. CYP genes that confer resistance to insecticides were then considered separately from CYP genes associated with the detoxification of other xenobiotics, such as those offering resistance to naturally occurring plant allelochemicals. We find a significant enrichment of TEs around CYP genes associated with resistance to synthetic insecticides in *P. nigronervosa*, *A. pisum*, *M. cerasi*, and *M. persicae* (Figure 7B). With the exception of *P. nigronervosa*, these are all species of “*most agricultural importance*” (Van Emden and Harrington 2017), and are therefore likely to have experienced particularly high exposure to synthetic insecticides, and extremely strong selection to evolve resistance. The significant enrichment of TEs at XGFs associated with resistance to synthetic insecticides in these species is thus consistent with their selective recruitment for resistance evolution. However, no significant enrichment of TEs around CYP genes associated with resistance to synthetic insecticides was found in 8 other species of “*most agricultural importance*” included in our analysis: *A. craccivora, A. gossypii, D. noxia*, *M. dirhodum, R. padi, S. graminum,* and *S. avenae* (Van Emden and Harrington 2017).

### Evidence for the selective enrichment of TEs at XGFs

We hypothesised that TE enrichment at resistance loci could arise through two distinct processes: (i) Chromatin availability, whereby XGFs are surrounded by open chromatin due to frequent transcriptional activity or constitutive expression, and so represent locations that are more accessible for transposition; (ii) Selective retention of beneficial TE insertions. Given that XGFs are essential for the detoxification of harmful plant metabolites and insecticides, and so are of key importance for survival and reproduction, TE insertions that modify XGFs in beneficial ways may be selectively retained and so spread through host populations. Consequently, the combined effects of XGFs as selective hotspots (especially considering the extremely strong selection mediated by intensive insecticide regimes), and the demonstrated evolutionary potential of TEs, could result in the selective retention of TE insertions surrounding XGFs over time. Such effects may be magnified by the influence of host stress under insecticide exposure, which has been implicated in increases in TE activity (Chénais et al. 2012; Stapley et al. 2015; Horváth et al. 2017). Specifically, increased TE activity may result in a higher likelihood of TE insertions occurring at any genomic location, including XGFs. However, TE insertions at XGFs may be more likely to be selectively retained, if they offer host benefits in the form of resistance mutations, due to their capacity to contribute to host evolvability through diverse genetic mechanisms (Bourque et al. 2018; Rebollo et al. 2012).

Only germline TE activity, as opposed to somatic TE activity, is relevant from the perspective of TE accumulation at XGFs, since only TE insertions that occur in the germline have the potential to be transmitted to the next generation (novel TE insertions in the genomes of somatic cells are an evolutionary dead-end). TE density at a gene, defined as the proportion of its sequence occupied by TEs, is positively correlated with germline gene expression, suggesting that TEs preferentially insert into regions that are actively transcribed in the germline (McVicker and Green 2010). Therefore, a key question to distinguish between alternative hypotheses that explain TE enrichment at XGFs is, to what extent are XGFs expressed in the germline? We addressed this question using RNA-seq data from a recent study on *A. pisum* (Jaquiéry et al. 2022) to examine gene expression in testes and ovaries for XGFs and a large panel of housekeeping genes. Housekeeping genes are constitutively expressed in all cell types and so represent a ‘maximally accessible’ case with which to examine TE enrichment (Butte et al. 2001) (Supplemental Table S4). We find that germline housekeeping gene expression is significantly higher compared to XGFs considering mean RPKM (reads per kilobase of transcript per million base pairs sequenced) in *A. pisum* (Wilcoxon Rank Sum, W_612,112_ = 52252, p < 0.01, Figure 8C). The observed pattern of expression of housekeeping genes and XGFs is also present in *Drosophila melanogaster*, suggesting conservation across insect diversity (Supplemental Figure S4). Consistent with chromatin availability as an explanation for TE enrichment, we find that TEs are significantly enriched at housekeeping genes compared to all other host genes, excluding XGFs (TE coverage: Wilcoxon Rank Sum, W_546,475492_ = 218,813,232, p < 0.01; TE count: Wilcoxon Rank Sum, W_546,475492_ = 170,683,160, p < 0.01), to a similar level as that observed for XGFs versus all other host genes (Figure 8A-B, Supplemental Figure S5). However, given that germline expression of XGFs is consistently low across all XGF types (Figure 8C), we conclude that germline expression cannot explain the observed enrichment of TEs at XGFs. Instead, our findings support the alternative explanation for the enrichment of TEs at XGFs, which is selective retention of TE sequences due to genetic contributions towards the evolution of xenobiotic resistance mutations. There is no mechanistic basis to suggest that TEs specifically target the regions surrounding XGFs as insertion sites, compared to other genomic locations. Instead, under this model TE insertions at XGFs are more likely to be retained and spread through the host population (compared to TE insertions at other loci), due to the combination of their capacity to contribute to host evolvability, and the extremely strong selection pressure imposed by insecticide treatment.

**Figure 8.**
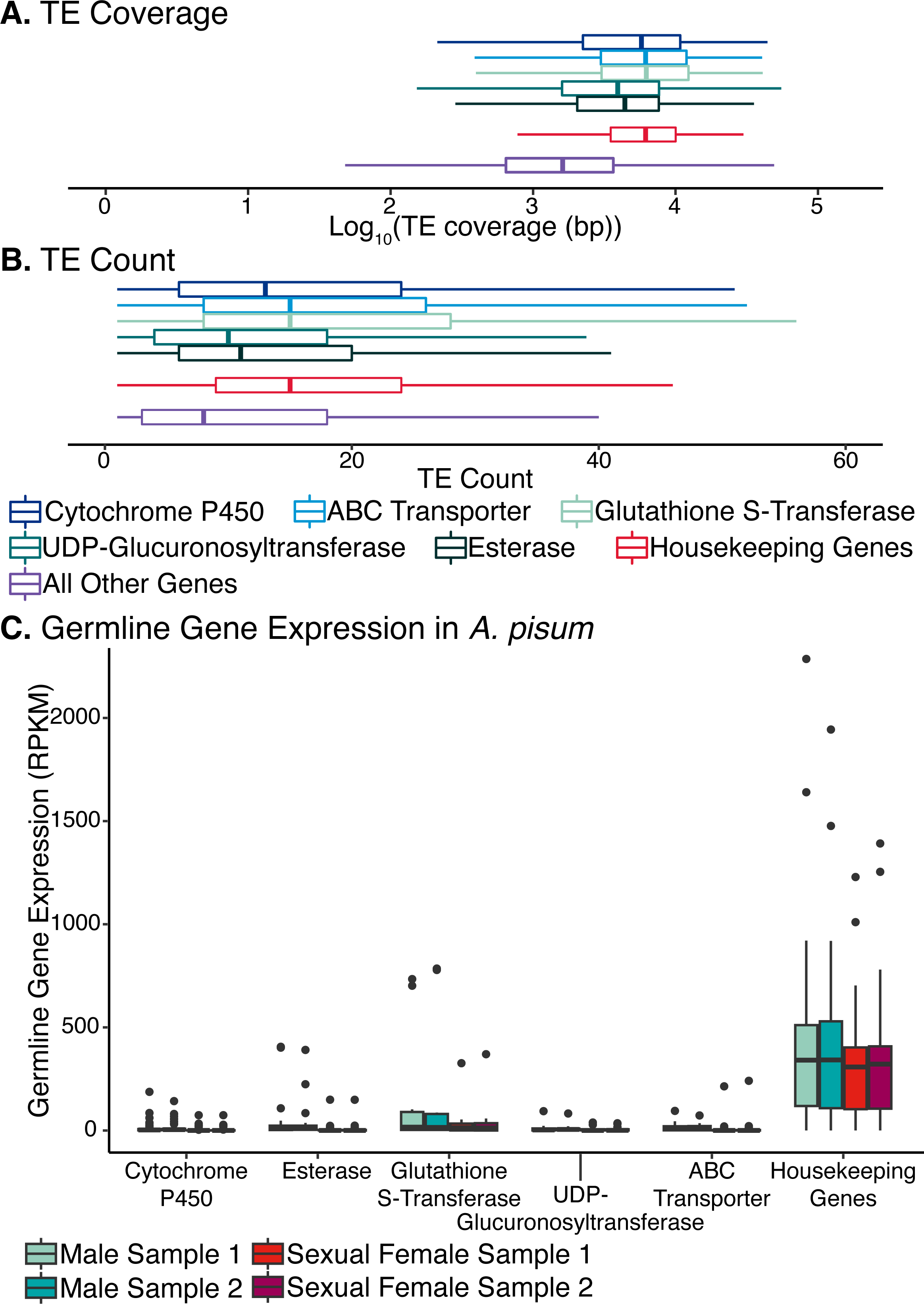
TE occupancy at XGFs, housekeeping genes, and all other genes, and expression of XGFs and housekeeping genes in A. pisum (Jaquiéry et al. 2022). A. TE coverage around XGFs, housekeeping genes, and all other genes. Box limits indicate upper and lower quartiles, whilst central lines indicate mean TE coverage. B. TE count around XGFs, housekeeping genes, and all other genes. Box limits indicate upper and lower quartiles, whilst central lines indicate mean TE count. C. Expression of XGFs and housekeeping genes in germline tissues (testis and ovary) in A. pisum, expressed as reads per kilobase of transcript per million base pairs sequenced (RPKM), to determine expected expression of XGFs in germline cell types. Gene types are indicated by colours in the key.

## Discussion

Resistance evolution remains a major societal challenge that greatly affects efforts to control harmful pests and pathogens (Van Emden and Harrington 2017). To develop new approaches to overcome the challenge of resistance evolution, a deeper understanding of the genomic mechanisms leading to the repeated emergence of resistant phenotypes is required. Whilst isolated examples of TE involvement in the evolution of xenobiotic resistance exist, the extent to which TEs act as a source of genomic novelty for resistance evolution, and host evolution more generally, remains an open question.

We examine whether genes implicated in xenobiotic resistance are enriched for TE insertions, potentially due to the generation of novel advantageous mutations for resistance evolution. We find an enrichment of TEs, both in terms of TE sequence coverage and number of TE insertions, at XGFs across all aphids and all XGF types. Further, we provide evidence that this enrichment is the consequence of selection at XGFs, as opposed to elevated insertion rates due to chromatin availability at XGFs in the germline.

Our results also uncover considerable variability in the enrichment of TEs around XGFs both within and among aphid species and XGF types. This demonstrates that TEs are not globally enriched at every XGF, but instead are enriched only at certain XGFs and depleted at others. Consequently, the patterns we observe are driven by hotspots of TE enrichment around certain XGF types, and in some cases individual XGFs, which show extremely high levels of enrichment compared to other XGFs. For example, considering all aphid species examined, we identify significant enrichment of TEs around GSTs. However, this result is driven by a subset of aphids, with 11 of the 20 species showing no evidence of significant enrichment around this XGF class (Supplemental Figure S4). This is consistent with varying levels of selection acting on particular detoxification genes and particular aphid species. Meanwhile, we find markedly more TE hotspots at UGTs and CYP genes, whereby certain XGFs are massively enriched for TE fragments. These observations demonstrate the nuances involved in patterns of TE accumulation at XGFs and underline a need for functional validation analyses to test the impact of individual TE insertions on XGF expression, and whether highly TE-enriched loci are indeed involved in xenobiotic resistance.

Important crop pests are likely to experience much higher levels of insecticide stress than non-target species, demonstrated by the increased number and diversity of insecticide resistance mutations in target species, compared to non-target species (Mota-Sanchez and Wise 2023). Consequently, variability in TE enrichment is expected among aphid species and XGFs, depending on variation in the degree of insecticide exposure. This prediction is confirmed when considering CYP genes, for which data clarifying the relationship to specific insecticides is available (rather than just to xenobiotics in general). Specifically, we find that TE enrichment is significantly higher around CYP genes that are implicated in the detoxification of synthetic insecticides compared to those involved in the detoxification of natural xenobiotics, but only in three out of ten aphids of “*major agricultural importance*” (Van Emden and Harrington 2017) included in our study (*A. pisum, M. cerasi*, and *M. persicae*). Consequently, either CYP genes are less important for survival in the other species (perhaps due to differences in pest control strategies, such as a greater reliance on forms of non-insecticide-based control, such as biocontrol strategies), or there are genomic mechanisms that limit the co-option of TEs for resistance evolution in these species.

Demonstrated contributions of TEs to xenobiotic resistance often involve small fragments of the original TE insertions, and contain motifs that act as host regulatory elements (Singh et al. 2020; Remnant et al. 2013; Panini et al. 2021; Bogwitz et al. 2005; Itokawa et al. 2011; Schmidt et al. 2010; Darboux et al. 2007; Gahan et al. 2001; Yang et al. 2007). Meanwhile, the emergence of a resistant phenotype frequently appears to result from a single, or up to a few, TE insertions at each specific XGF locus, rather than large accumulations of TE DNA (Schmidt et al. 2010; Daborn et al. 2002; Chung et al. 2007; Panini et al. 2021). This presumably occurs due to genomic and selective processes that act to retain the core TE regions that convey a selective benefit, while purging long repetitive and potentially deleterious regions (e.g. that promote non-homologous recombination with similar repetitive elements). Meanwhile, the impacts of these TE insertions can occur via complex multistep processes including the retention of several truncated TE fragments (such as for *CYP6G1*, Schmidt *et al*. (2010)). Thus, a key consideration is that TE-associated resistance mechanisms go beyond simple enrichment at specific loci.

We find that DNA TEs are highly enriched at XGFs, despite the more widely acknowledged role of TEs with more complex *cis*-regulatory elements, such as LTR retrotransposons, in altering host gene expression (Chung et al. 2007). This may suggest that other processes beyond TE-mediated increases in expression are involved in the patterns of enrichment we uncover. Although, it is entirely possible that the DNA TEs involved also contain sequences relevant for the recruitment of transcription factors that increase expression. Meanwhile, at present we cannot currently explain the repeated cases whereby certain XGFs are massively enriched in terms of TE coverage and TE count compared to other XGFs. Thus, unravelling the influence of TEs at these loci represents a key target for future research.

Gaining a deeper understanding of the mechanisms that drive patterns of association between TEs and XGFs will require contributions from complementary approaches to the broadscale comparative genomic analyses performed here. Of particular relevance are functional validation studies to examine the influence of individual TE insertions on nearby XGF loci, which our results provide considerable scope to address. Further analyses are also required to elucidate the underlying mechanisms by which TE sequences may contribute to resistance phenotypes, such as screening to investigate if implicated TE sequences contain transcription factor binding sites, as well as information on chromatin conformation and the availability of DNA within TE sequences at XGFs. Meanwhile, detailed intra-species analyses can provide valuable evolutionary insights into individual TE insertions of interest (e.g. Aminetzach et al. (2005)), and information on TE occupancy within populations (e.g. Barrón et al. (2014)). Future work considering strains displaying different resistance phenotypes using high-quality long-read datasets and individual TE annotations (e.g Rech et al. (2022)) will enable the characterisation of TE variants likely contributing to resistance evolution in aphids. Collectively, such approaches will ultimately allow evaluation of the extent to which TEs are utilised in resistance mutations, versus the contributions arising from other classes of genetic variation such as SNPs and CNVs.

## Materials and Methods

### Estimating Host Phylogeny

The genome assemblies of 20 aphid species, and 7 diverse hemipteran outgroup species (*Ericerus pela*, *Maconellicoccus hirsutus*, *Phenacoccus solenopsis, Bemisa tabaci, Sogatella furcifera, Diaphorina citri, Pachypsylla venusta*), were obtained along with gene annotations where available (Supplemental Table S6). To estimate host phylogeny, a supermatrix approach was used. Specifically, for each genome assembly, BUSCO (version 5.2.2) (Manni et al. 2021a, 2021b) was used with the *hemiptera_odb10* geneset to identify conserved gene orthologs in each genome assembly. The identifiers for all complete genes were extracted and those present in less than three genomes were removed. The amino acid sequences for each gene from all species were extracted and saved in individual FASTA files. For each gene, sequences were aligned using MAFFT (version 7.453) (Katoh and Standley 2013) (-- auto). The subsequent alignments were concatenated using Phykit create_concat (Steenwyk et al. 2021) to generate a supermatrix. To generate the species phylogeny, we performed 1,000 ultrafast bootstrap repetitions (-bb 1000) in IQ-TREE (version 1.6.12) (Nguyen et al. 2015), using the best fit amino acid model identified by Modelfinder (Kalyaanamoorthy et al. 2017) (Supplemental File S1).

### Transposable Element Annotation

To curate and annotate TEs, each genome assembly was annotated with the Earl Grey TE annotation pipeline (version 1.3) (https://github.com/TobyBaril/EarlGrey) (Baril et al. 2022). Briefly, known TEs from Arthropoda (-r arthropoda) were first annotated using both the Dfam 3.4 (Hubley et al. 2016) and Repbase RepeatMasker Edition (version 20181026) (Jurka et al. 2005; Kapitonov and Jurka 2008) TE databases. Following this, Earl Grey identified *de novo* TE families and refined these using an iterative ‘BLAST, Extract, Extend’ process (Platt et al. 2016). Following final annotation with the combined library of known and *de novo* TE families, annotations are processed to remove overlapping annotations and to defragment annotations likely to originate from the same TE insertion. TE annotation GFF files are provided in Supplemental File S2.

To characterise TEs as either shared or species-specific, *de novo* TE libraries from each genome assembly were clustered to the TE family definition of Wicker et al. (2007), implemented as described by Goubert et al. (2022) using cd-hit-est (-d 0 -aS 0.8 -c 0.8 -G 0 - g 1 -b 500 -r 1) (Li and Godzik 2006; Fu et al. 2012). Sequences within each cluster were designated a number between 0 and 20 to determine the number of other species that the TE family was shared with, with 0 identifying TEs unique to a species, and 19 identifying TEs shared across all 20 aphid species considered in this study. For known TE families from Dfam and Repbase, each TE was labelled using the same system based on the number of species the TE family was identified in.

Genetic distance from TE consensus sequence is used as a proxy for estimated timing of TE activity (i.e TE age). Using genetic distance from TE consensus as a proxy for insertion time should be taken with caution, as this can be influenced by the methodology used to generate TE consensus sequences, and whether known TE sequences from other organisms are used as a reference or purely *de novo* TE consensus sequences from the species of interest. However, this metric can still be used to provide a rough relative estimate of TE activity.

### Phylogenetic Heritability of TE Abundance and Diversity

To determine the phylogenetic heritability of TE abundance, expressed as total TE count, and TE diversity, expressed as total number of distinct TE families, we measured the amount of variation explained by phylogenetic relationships using Bayesian phylogenetic mixed models with Markov chain Monte Carlo estimations, with an intercept fitted as a fixed effect and the phylogeny fitted as a random effect, in the MCMCglmm package (Hadfield 2010) in R (version 4.2.1) (R Core Team 2023) using the Rstudio IDE (RStudio Team 2015; Racine 2013). A poisson error distribution and log link function were used, and inverse gamma priors were specified for all R and G-side random effects (V = 1, ν = 0.002). Models were run for 11,000,000 iterations with a burn-in of 1,000,000 and a thinning interval of 1,000. This approach generated 10,000 posterior samples from which posterior mode and 95% CIs were calculated. The proportion of between-species variation explained by phylogeny was calculated from the model using the equation *V_p_ / (V_p_ + V_s_)* where *V_p_* and *V_s_* represent the phylogenetic and species-specific components of between-species variance (Freckleton et al. 2002), which is equivalent to phylogenetic heritability.

### Gene Predictions and Transposable Element Association with Genomic Compartments

For genome assemblies lacking a gene annotation GFF file at the time of analysis (*A. gossypii*, *H. cornu, M. dirhodum, P. nigronervosa, R. padi, S. graminum, S. chinensis, S. avenae, S. miscanthi*), genes were annotated using AUGUSTUS (version 3.3.3) (Stanke et al. 2008; Keller et al. 2011) with the pea aphid reference gene set (--species=pea_aphid -- strand=both --genemodel=partial --introns=on --exonnames=on --gff3=on).

To prepare genome annotation files for intersection with TE loci, each gene annotation GFF was processed to generate a GFF file containing coordinates of exons, introns, 5’ and 3’ flanking regions, and intergenic space. Gene flanking regions are defined here as 20kb directly upstream or downstream of the gene body. These flanking regions are determined to identify TEs at a distance that could be described as the proximate promoter region, rather than just accounting for the core promoter region, to include both promoter and more distal enhancer regions for genes (Lenhard et al. 2012). Intergenic space coordinates were generated using BEDTools complement (Quinlan and Hall 2010), to generate inverse coordinates to those of genic compartments and flanking regions. To quantify TE occupation in different genomic compartments (introns, exons, 5’ and 3’ flanking regions, and intergenic space), BEDTools intersect was used to calculate overlap (-wao) between all compartments and TEs.

### Identification of Xenobiotic Gene Family Loci and Housekeeping Genes

The gene identities for all ABC transporters, cytochrome P450s, UDP-glucuronosyltransferases, esterases, and glutathione S-transferases were manually curated from the functional annotation of the *M. persicae* G006v2 genome (Singh et al. 2021). Using these identifiers, exon coordinates were obtained from the *M. persicae* gene annotation GFF3 file, and corresponding nucleotide sequences were extracted from the genome assembly. The resulting exon sequences from *M. persicae* were used as queries to identify XGFs in all other aphid assemblies using BLASTN (Camacho et al. 2009) with an e-value threshold of 1×10^-10^. BLAST hits for each species were manually inspected to extract those that best represented each XGF, with the inclusion of gene duplications where appropriate. Following this, a BED file of XGF coordinates for each species was generated including the annotation of flanking regions as described above (Supplemental File S3).

Housekeeping genes (HKGs) are required for basic cellular functions and are typically expressed in all cell types (Butte et al. 2001). Amino acid sequences of 28 HKGs characterised for aphids (Bansal et al. 2012; Koramutla et al. 2016; Li et al. 2020; Kang et al. 2017; Yang et al. 2015, 2014) were obtained from GenBank (Benson et al. 2013) for *M. persicae* (Supplemental File S4). Subsequently, a BLASTX search was performed, with default parameters, against each aphid genome to identify HKG loci in the other 19 species. For each species, HKGs were manually curated and a BED file of HKG coordinates, including the annotation of flanking regions, was generated (Supplemental File S5). To quantify TE occupation around XGFs and HKGs, BEDTools intersect was used to calculate overlap (-wao) between annotated TEs and XGFs.

## Acknowledgements

TB was supported by a studentship from the Biotechnology and Biological Sciences Research Council-funded South West Biosciences Doctoral Training Partnership (grant number BB/M009122/1).

AP was supported by the European Research Council (ERC) under the European Union’s Horizon 2020 research and innovation program (grant agreement number 646625).

CB was supported by a Biotechnology and Biological Sciences Research Council (BBSRC) grant (grant number BB/S006060/1) and the European Research Council (ERC) under the European Union’s Horizon 2020 research and innovation program (grant agreement number 646625)

AH was supported by a Biotechnology and Biological Sciences Research Council (BBSRC) David Phillips Fellowship (grant number BB/N020146/1).

*Myzus cerasi* DNA sequencing data were downloaded from AphidBase. Funding for the sequencing was provided by ERC Starting Grant APHIDHOST-310190 awarded to Jorunn Bos at the James Hutton Institute, United Kingdom.

For the purpose of open access, the author has applied a ‘Creative Commons Attribution (CC BY) licence to any Author Accepted Manuscript version arising.

## Author Contributions

TB performed TE annotation and downstream analyses, produced the figures, and drafted the manuscript. AP manually curated xenobiotic resistance genes from *M. persicae.* CB and AH contributed to the analytical design and data analysis, and participated in writing the manuscript. AH conceived and coordinated the study. All authors read and approved the final manuscript.

