## Supplementary material for "Transposon accumulation at xenobiotic gene family loci: a comparative genomic analysis in aphids": Sup_Fig_S1

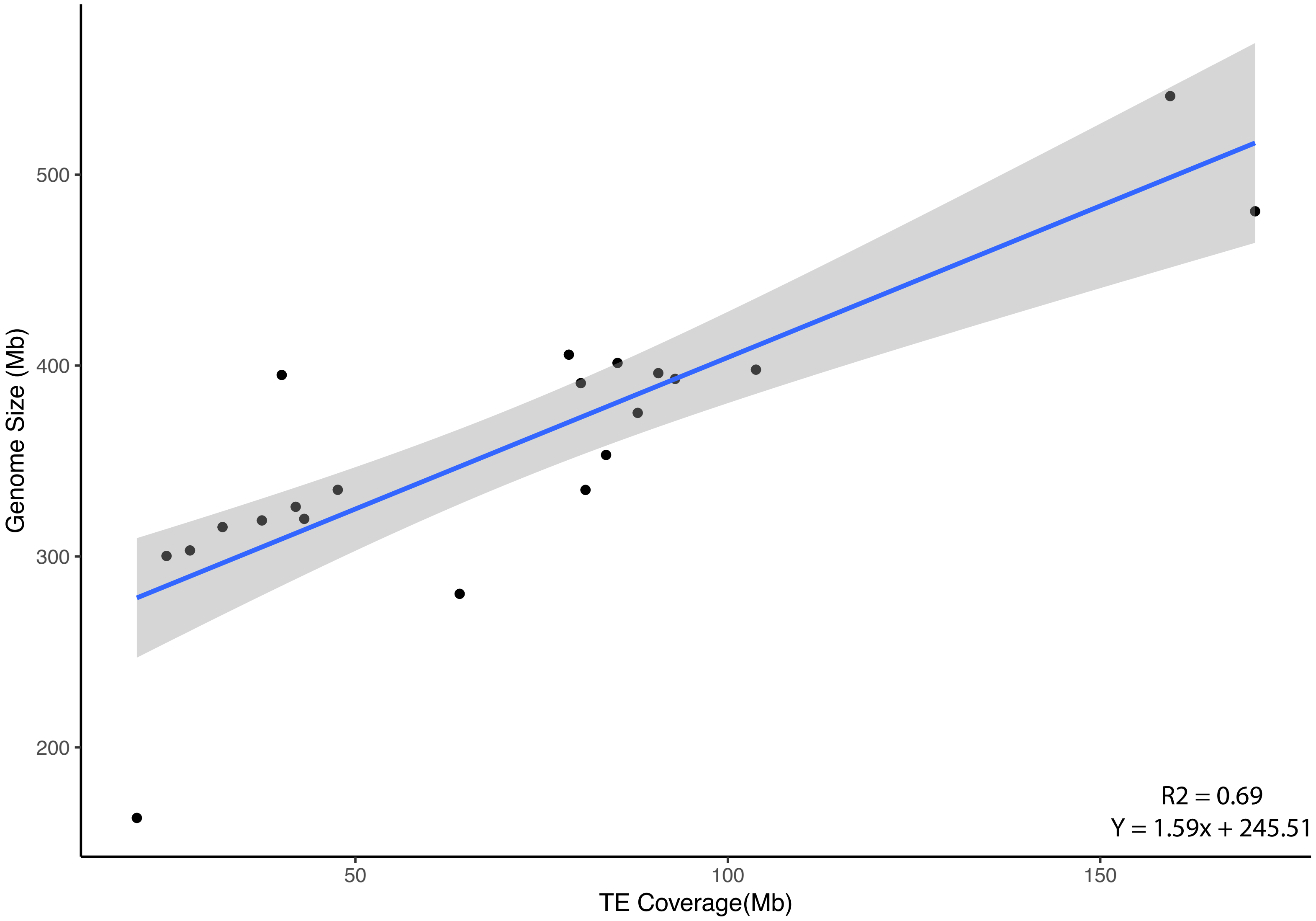


Supplemental Figure S1. The relationship between TE content and genome size in aphids.
Blue line indicates the relationship calculated using linear regression. Grey area indicates 95% confidence intervals.
