## Supplementary material for "Transposon accumulation at xenobiotic gene family loci: a comparative genomic analysis in aphids": Sup_Fig_S2

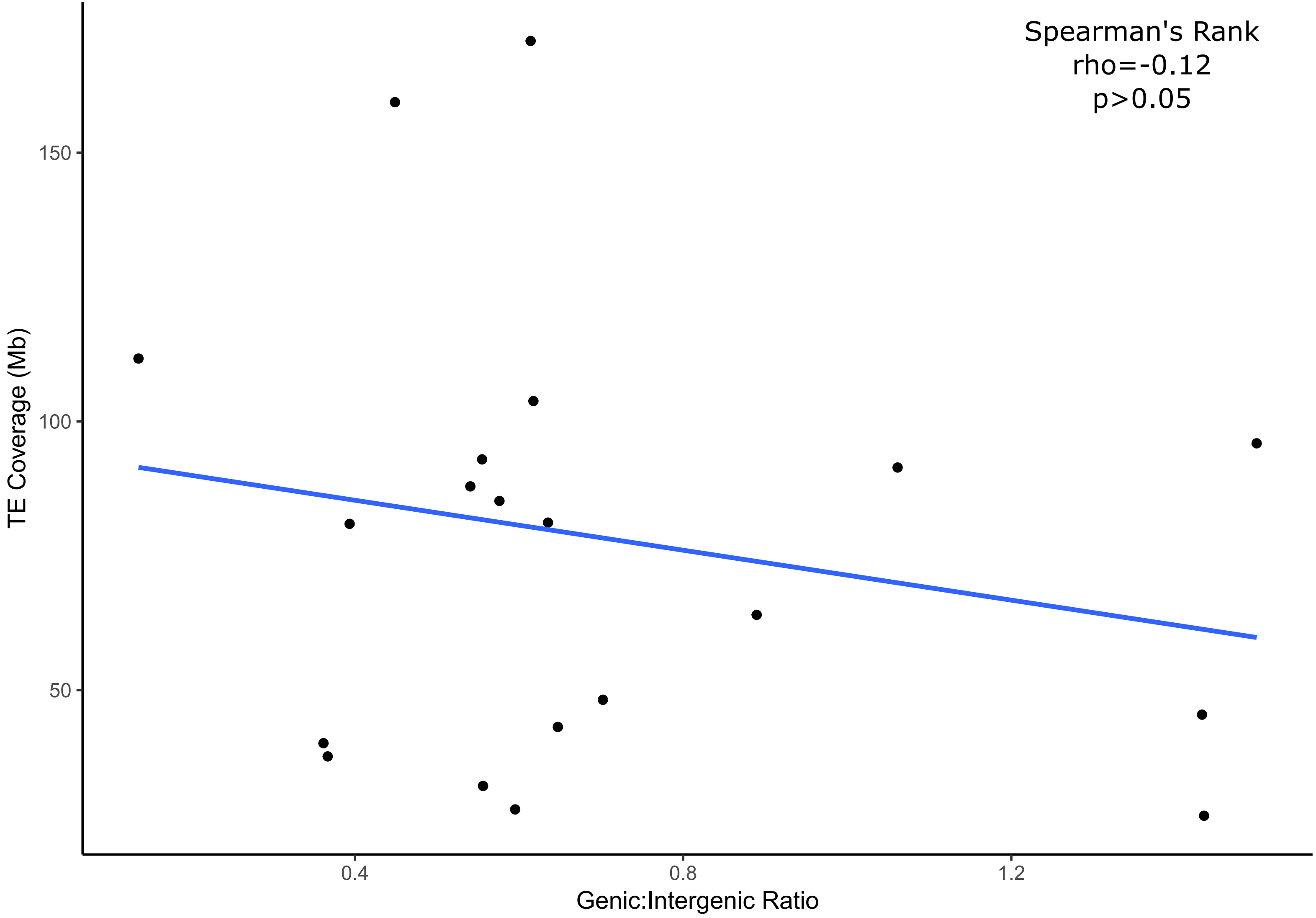


Supplemental Figure S2. Correlation between genic:intergenic base pairs ratio and TE content in aphids.
