## Supplementary material for "Transposon accumulation at xenobiotic gene family loci: a comparative genomic analysis in aphids": Sup_Fig_S3

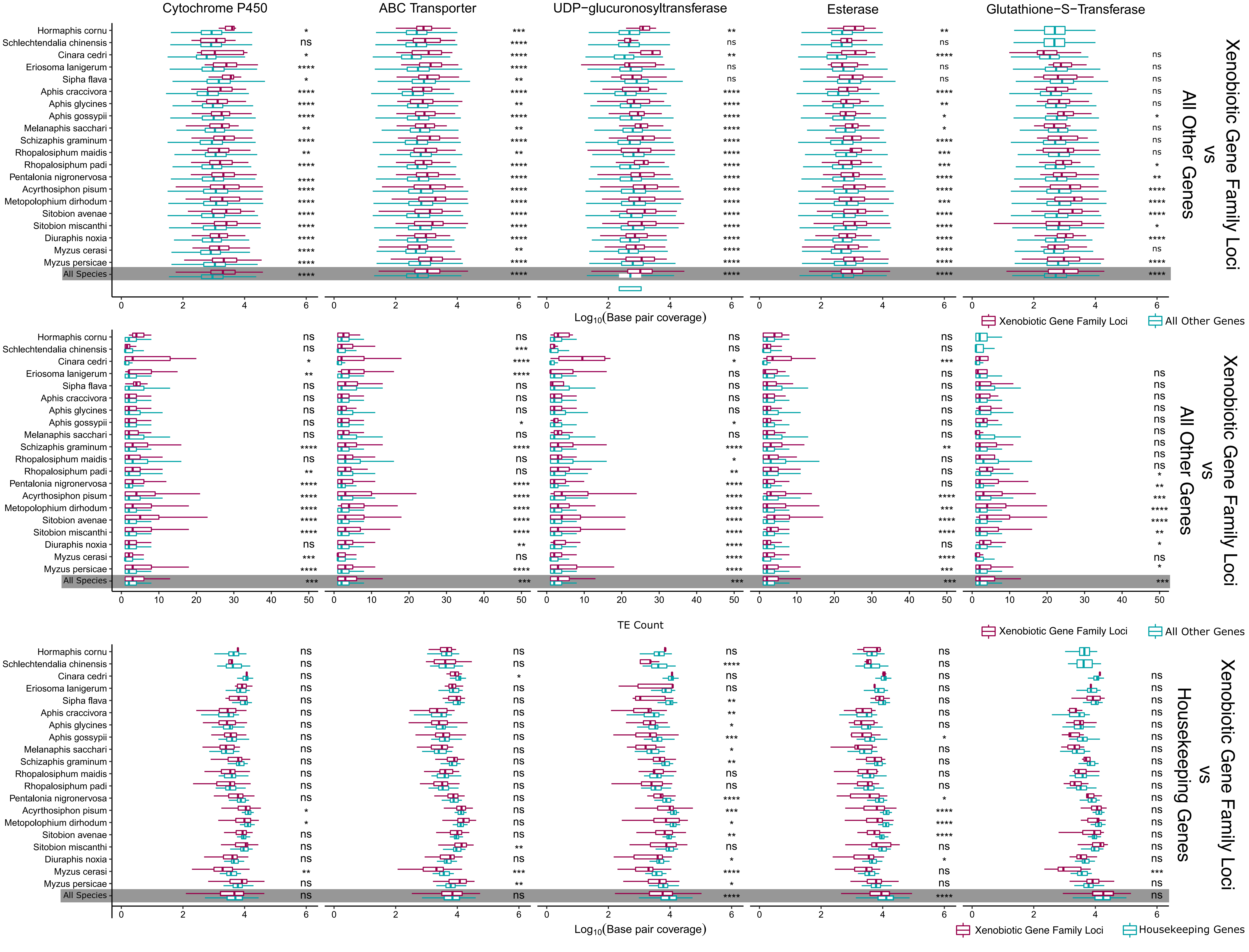


Supplemental Figure S3. TE abundance around xenobiotic gene family loci compared to all other genes and housekeeping genes, split by xenobiotic gene family type.
