## Supplementary material for "Transposon accumulation at xenobiotic gene family loci: a comparative genomic analysis in aphids": Sup_Fig_S4

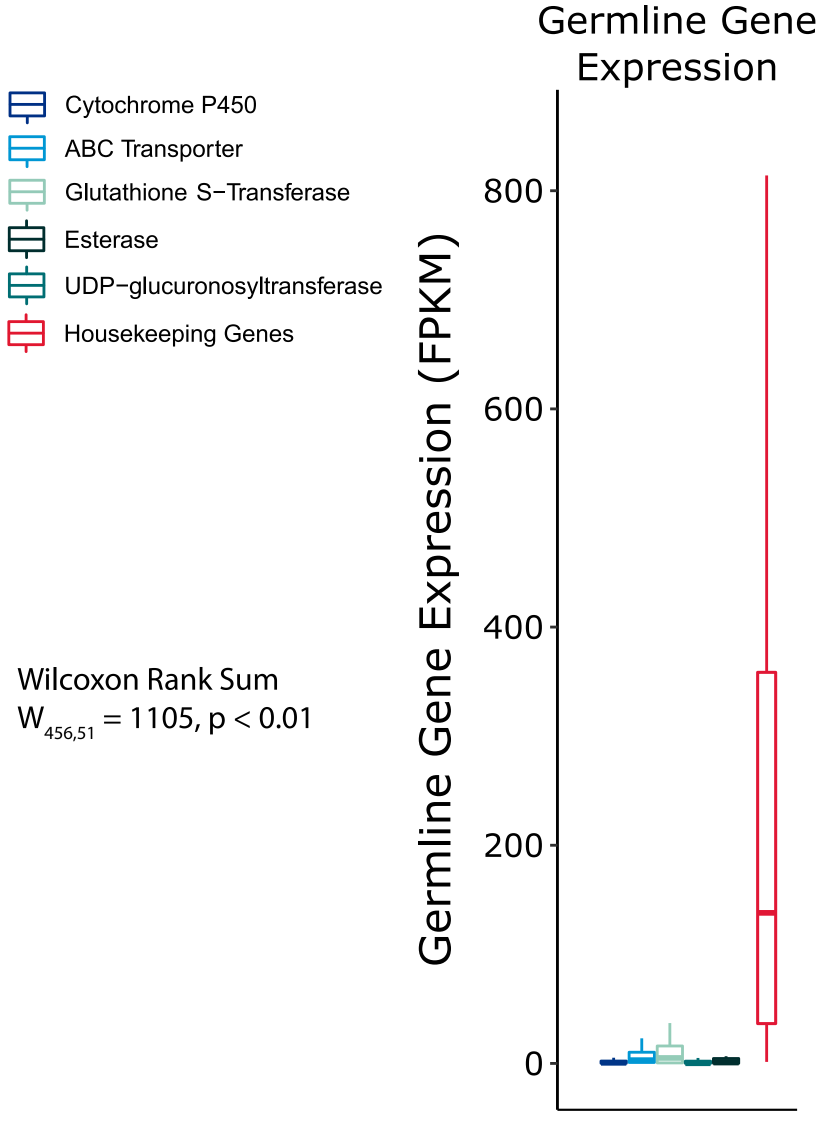


Supplemental Figure S4. Expression of XGFs and housekeeping genes in germline tissues (testis and ovary) in D. melanogaster using data from FlyAtlas2 (Krause et al. 2022). Expression is measured as fragments per kilobase of transcript per million base pairs sequenced (FPKM), to determine expected expression of XGFs in germline cell types. Gene types are indicated by colours in the key.
