## Supplementary material for "Transposon accumulation at xenobiotic gene family loci: a comparative genomic analysis in aphids": Sup_Fig_S5

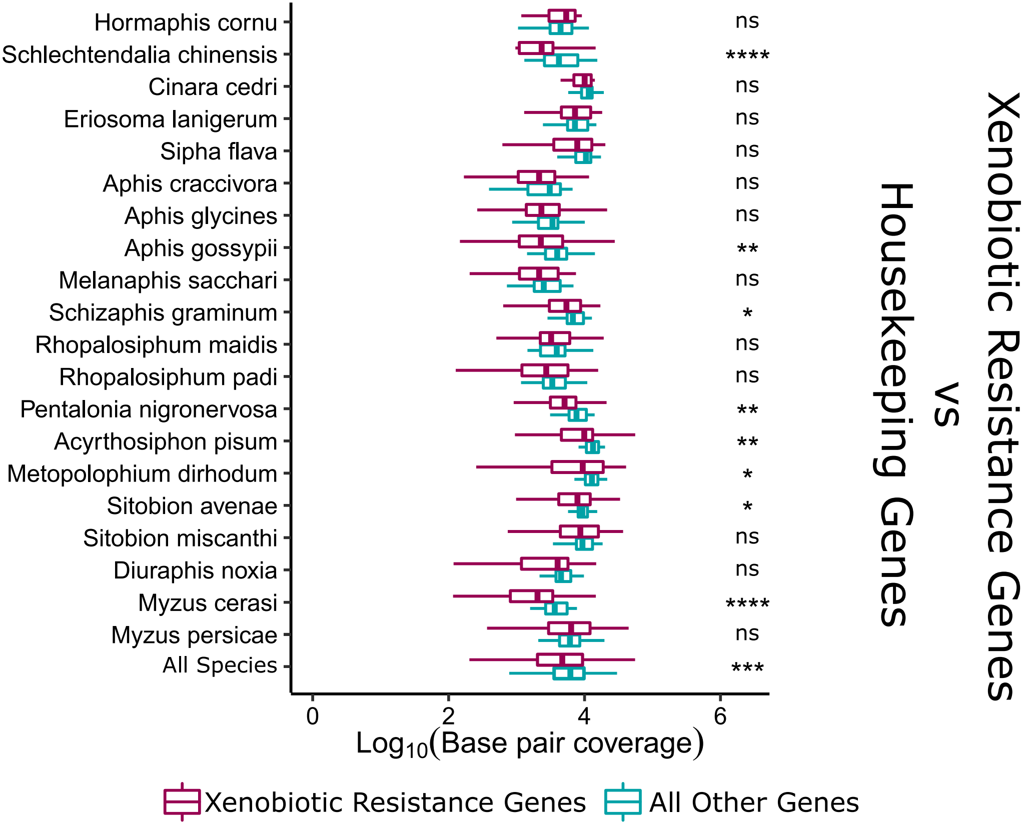


Supplemental Figure S5. TE coverage around xenobiotic gene family loci compared to housekeeping genes.
